# The history and evolution of the Denisovan-*EPAS1* haplotype in Tibetans

**DOI:** 10.1101/2020.10.01.323113

**Authors:** Xinjun Zhang, Kelsey Witt, Amy Ko, Kai Yuan, Shuhua Xu, Rasmus Nielsen, Emilia Huerta-Sanchez

## Abstract

Recent studies suggest that admixture with archaic hominins played an important role in facilitating biological adaptations to new environments. For example, interbreeding with Denisovans facilitated the adaptation to high altitude environments on the Tibetan Plateau. Specifically, the *EPAS1* gene, a transcription factor that regulates the response to hypoxia, exhibits strong signatures of both positive selection and introgression from Denisovans in Tibetan individuals. Interestingly, despite being geographically closer to the Denisova cave, East Asian populations do not harbor as much Denisovan ancestry as populations from Melanesia. Recently, two studies have suggested two independent waves of Denisovan admixture into East Asians, one of which is shared with South Asians and Oceanians. Here we leverage data from *EPAS1* in 78 Tibetan individuals to interrogate which of these two introgression events introduced the *EPAS1* beneficial sequence into the ancestral population of Tibetans, and we use the distribution of introgressed segment lengths at this locus to infer the timing of the introgression and selection event. We find that the introgression event unique to East Asians most likely introduced the beneficial haplotype into the ancestral population of Tibetans around 43,000 (15,700–60,000) years ago, and selection started 12,000 (1,925-50,000) years ago. Our estimates suggest that one of the most convincing examples of adaptive introgression is in fact selection acting on standing archaic variation.

## Introduction

The identification of the Denisovan genome using DNA recovered from a phalanx bone is one of the most stunning discoveries in human evolution in the past decade^1,2^. However, many questions remain unanswered regarding the Denisovans. For example: What did the they look like? What was their geographical range? What is their genetic legacy to modern humans? Much of the ongoing research investigating the Denisovans focuses on studying the morphological features from dental and cranial samples^3^, dating the age of remains from the Denisova cave^4^, and learning about the admixture events that involved Denisovans, Neanderthals, and other unknown archaic populations^1,5–7^. We now know that Denisovans diverged from Neanderthals approximately 390 thousand years ago (ka)^8,9^, and both groups inhabited Eurasia until up to 40 ka^4,10^ based on radiocarbon dating of materials from Neanderthal or Denisovan archeological sites.

Although the fossil remains of Denisovans found so far are limited in number and highly fragmented in nature^1,11,12^, certain aspects of this hominin group have been revealed through studying a single high-coverage genome^2^. The occurrence of admixture between archaic hominins and modern humans is undisputed, as it left varying amounts of archaic DNA in our genomes at detectable levels^1,8,13^. Notably, Papuans and indigenous Australians harbor the largest genome-wide amount of Denisovan introgression (~ 1% - 5%^1,6,14,15^), followed by East and South Asians (~ 0.06 % - 0.5%^1,6,14^), and Indigenous Americans (~ 0.05% - 0.4%^1,6,14^). Thus, one approach to study the Denisovans is through the surviving Denisovan DNA segments in modern humans.

Examination of Denisovan-like DNA in modern humans revealed a number of candidate genes with robust signatures of adaptive introgression^16–21^, among which the most well-known example is found in the *Endothelial Pas Domain Protein1* gene (*EPAS1*) in modern Tibetans^22–24^ that facilitated local adaptation to their high altitude and hypoxic environment. The discovery of adaptive introgression in Tibetans is particularly striking, as they do not carry high amounts of Denisovan ancestry genome-wide, compared to other South Asian and Oceanian populations^6^. Conversely, the Oceanian populations - including the Papuans - do not carry the Denisovan *EPAS1*-haplotype, most likely due to the absence of selective pressure outside of the high-altitude environment. The Tibetan plateau, with an average altitude above 3,500 meters and oxygen concentration considerably lower than at sea level, creates a strong physiological stress for most humans. One common acclimatization to the hypoxic environment is an increase in hemoglobin concentration^25^, which increases blood viscosity and is associated with increased risk of pregnancy complications and cardiovascular disease^26,27^. Remarkably, Tibetans have a severely blunted acclimatization response compared to lowlanders at high altitudes and tend not to suffer from clinically elevated hemoglobin concentration^28^. This presumed adaptive response is directly associated with variants in the *EPAS1* gene, which encodes a transcription factor in the hypoxia response pathway.

The remarkable Denisovan connection to Tibetans’ high-altitude adaptation has led to more questions regarding this already mysterious hominin group. For example, why are populations with Denisovan ancestry, including the Tibetans and Oceanians, located far away from the Denisova cave in Siberia? One explanation for these seemingly puzzling findings is a large Denisovan geographical range. Multiple introgression events may explain why some human populations exhibit higher levels of Denisovan introgression despite being located far away from the Altai mountains in Siberia. Indeed, Browning *et al*.^7^ proposed two Denisovan introgressions into modern East Asians, one of which is shared with Papuans and South Asians. More recently, Jacobs *et al*.^29^ proposed an additional introgression event into the ancestral population of Papuans, making a total of three Denisovan introgression pulses in Asia. Their estimates of divergence between the three Denisovan groups that admixed with modern humans are large enough (~280 ka-360 ka) to suggest that there were multiple Denisovan-like hominin groups inhabiting diverse locations in Asia.

In this study, we investigate the surviving Denisovan introgressed segments in Tibetans to address the following questions: Do Tibetans exhibit signatures of more than one Denisovan introgression? If so, which introgression event introduced the beneficial *EPAS1* haplotype, and when? Did selection act immediately after introgression, or plausibly later when modern humans began inhabiting the Tibetan Plateau? To address these questions, we examined the *EPAS1* gene sequences from a combined dataset of 78 Tibetan individuals from two previously published studies^23,30^, among which 38 are high-coverage whole genome sequences^30^. We leveraged information from the introgressed tracts in Tibetans to infer the key time points related to the Denisovan introgression, as well as the onset of selection. We also employed the whole genomes in the combined dataset to demonstrate that the ancestors of modern Tibetans, similar to other East Asian populations^7^, experienced two Denisovan introgression events. Our results provide resolution to the East Asian-specific Denisovan admixture event that led to one of the most fascinating stories of human adaptation, and shed light on the effects of different evolutionary processes that shape patterns of adaptive introgression in humans.

## Results

### Evidence of three distinct archaic introgression episodes with ancestral Tibetans: one from Neanderthals and two from Denisovans

To characterize the genomic landscape of archaic introgression in Tibetans and to determine the number introgression pulses, we applied the method developed in Browning *et al*.^7^, SPrime. This is a reference-free method that detects sets of diagnostic SNPs that tag putatively archaic-introgressed segments in different regions of the genome. Applying SPrime to the autosomes of 38 Tibetan genomes^30^ we inferred 1,426 regions, each containing a set of diagnostic, putatively archaic-introgressed SNPs using Africans (YRI)^31^ as an outgroup. The remnants of archaic introgression in Tibetans are spread widely across the genome, illustrated by the presence of SPrime-inferred segments on all 22 autosomes (Supplementary Figure 1). Following Browning *et al*.^7^, for each segment, we computed the match rate of Tibetans against the Altai Neanderthal and Denisovan genomes at the different sets of diagnostic SNPs. The match rate distribution is visualized as a contour plot in Figure 1. Most regions detected show high affinity to Neanderthal (~80% matching) and low affinity to Denisovan representing the highest peak (colored in red) in the plot. This is consistent with the observation of higher rate of Neanderthal introgression compared to Denisovan introgression in all Eurasian populations^13^. We also observe two additional peaks representing segments of the genome that have low (~ 10%) affinity with Neanderthals and higher (~50% and ~80%) affinity with Denisovans. These two peaks represent putative Denisovan introgressed segments, and the bimodal distribution of Denisovan match rates is concordant with the hypothesis of two pulses of admixture with Denisovan-like archaic humans in East Asia^7^, as only one peak in the match rate (with the Altai Denisovan) distribution is expected under a single pulse of introgression.

Near *EPAS1*, two putatively archaic introgressed segments are inferred within 200kb up- and downstream of the *EPAS1* gene region, with a match rate to the Altai Denisovan of 72% and 46% respectively (Table 1, Supplementary Figure 2a). The previously identified segment within the *EPAS1* gene^23^ that harbors the adaptive allele was not detected by SPrime using YRI as the outgroup population. This is likely due to the fact that Yorubans carry a small number of the archaic alleles in the *EPAS1* region^23,32^, which could occur from mechanisms such as shared ancestry with archaic humans, unknown archaic admixture in Africa^33–35^, or backward gene flow from non-Africans to Africans^36^. Yorubans, however, do not carry the Denisovan haplotype at the *EPAS1* gene (Supplementary Figure 3), but the presence of a few archaic alleles in this region hinders the detection of introgressed segments using algorithms such as SPrime. In fact, repeating the SPrime analysis using modern Europeans (CEU, who do not harbor the Denisovan variants at *EPAS1*) as an outgroup population does detect the putatively adaptive archaic segment in the core region of *EPAS1* (chr2:46,550,132-46,600,661, hg19) that matches with high affinity to the Altai Denisovan (82.14%) but not the Altai Neanderthal (28.57%, see Table 1). In this region, the SPrime-inferred variants in Tibetans are more similar to the Denisovan (Supplementary Figure 3), and as expected, exhibits high genetic differentiation between Tibetans and Han Chinese (as measured by *F_ST_*, see Supplementary Figure 4). Using Europeans (CEU) as an outgroup for detecting introgression in Tibetans results in a similar genome-wide distribution of match rates as observed earlier, but with fewer inferred segments in general, indicating that these are primarily a subset of the ones we obtained when using the YRI as an outgroup (Supplementary Figure 2b).

**Table 1:** Archaic introgression segments within 200kb of the *EPAS1* gene region inferred by SPrime. The SPrime program infers three introgressed segments at or within 200kb range of the *EPAS1* gene in Tibetans. One segment is within the gene (core region), another segment is upstream of *EPAS1* and the third segment is downstream of *EPAS1*. Table shows the match rates to the Altai Denisovan and Neanderthal, the length of each segment, the chromosome and position range, number of diagnostic SNPs detected by SPrime, and the outgroup used by SPrime. The match rate is defined as the proportion of alleles at the SPrime diagnostic SNPs that are present in the sequenced archaic individual.

|  | Upstream region | Core region | Downstream region |
| --- | --- | --- | --- |
| **Segment Length** | 140.9 kb | 50.5 kb | 150.9 kb |
| **Positions (hg19)** | chr2:46,317,587-46,458,516 | chr2:46,550,132-46,600,661 | chr2:46,657,114-46,808,047 |
| **Number of Archaic-Specific Alleles** | 63 | 29 | 81 |
| **Match Rate with Altai Neanderthal** | 13.6% | 28.57% | 22.53% |
| **Match Rate with Altai Denisovan** | 72.73% | 82.14% | 46.48% |
| **Reference Outgroup** | YRI, CEU | CEU | YRI, CEU |
| **Input Data** |  | 38 Tibetan whole genomes<sup>30</sup> |  |

### The East Asian-specific Denisovan introgression event introduced the beneficial EPAS1 haplotype to ancestral Tibetans

We showed in the previous section that similar to other East Asians^7,29^, Tibetans also display evidence of two Denisovan introgression events, with one being unique to East Asians. Next, we tried to determine which of these two admixture events introduced the beneficial haplotype in *EPAS1*. To do so, we compared the introgressed segments in the 38 Tibetans at *EPAS1* to the SPrime-inferred regions that exhibit the highest (> 60%) Denisovan match rate and a low (< 40%) Neanderthal match rate (peak from Figure 1, Supplementary Figure 5). These segments were likely introduced via an East-Asian specific introgression event with a Denisovan population more closely related to the Altai Denisovan; other populations (e.g. South Asians, Oceanians) lack introgressed segments with this level of affinity to the Altai Denisovan^7^.

Since SPrime does not infer the introgressed segments for each individual chromosome, we applied a Hidden Markov Model (HMM)^13,17,37^ to infer the Denisovan-introgressed tracts in each Tibetan haplotype (see Methods). We show that the introgressed segments in *EPAS1* exhibit high Denisovan affinity, as shown in Figure 1, and are of similar length as other segments with the highest Denisovan affinity (Supplementary Figure 6a-b). Segments in *EPAS1* are only outliers in terms of the tract frequency (Figure 2a), which is in concordance with the expectation of positive selection acting on this region. Based on these observations, we propose that the *EPAS1* haplotype in Tibetans was introduced through the pulse of East Asian-specific Denisovan introgression.

**Figure 1:**
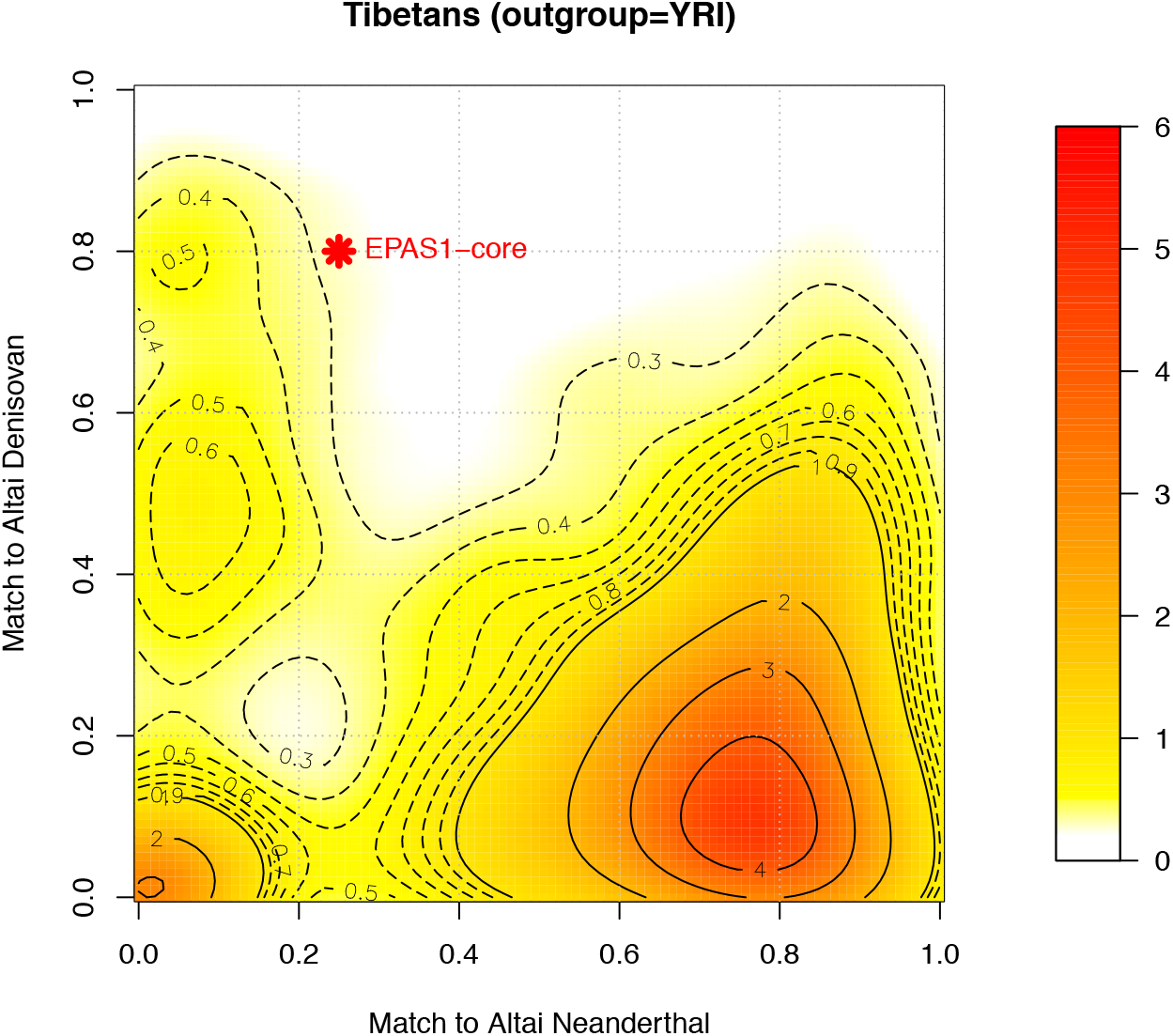
Introgressed segments in *EPAS1* and the genome-wide match rate with archaic individuals. This figure shows the density distribution of match rate to archaic individuals (Altai Denisovan or Neanderthal) in putatively archaic introgressed segments in 38 Tibetans, inferred by the SPrime program using Africans (YRI) as the outgroup. The match rate is defined as the proportion of alleles at the SPrime diagnostic polymorphic sites in a putatively introgressed segment that are present in the genome of archaic individuals at those positions ^7^. For a given segment, a match rate of 0 denotes that at a given set of diagnostic sites inferred by SPrime, none of the alleles at those sites match the corresponding alleles in the sequenced archaic human. The color range denotes the density of the contours, with red indicating high density and yellow indicating low density. The red star represents the matching coordinates for the introgressed segment within the *EPAS1* gene.

**Figure 2.**
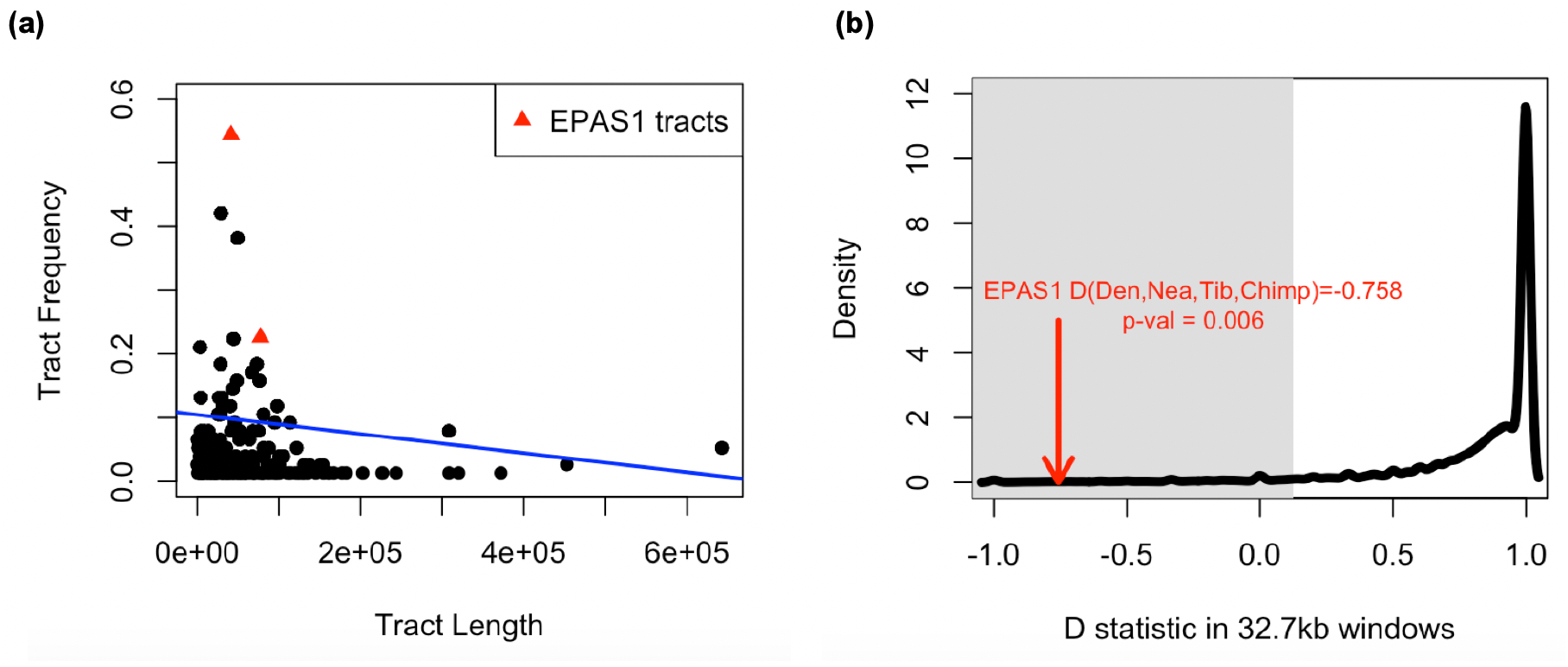
Introgressed tract length and frequency and D-statistics. Panel (a) shows the lengths (x-axis) and frequencies (y-axis) of introgressed tracts inferred by an HMM applied to the high Denisovan affinity regions detected with Sprime in 38 Tibetans. These regions have a match rate < 40% to Neanderthals and > 60% to Denisovans (see Figure 1 and Methods). The red triangles represent the introgressed tracts at *EPAS1*, including a long and a short segment (80kb and 40kb respectively). The tract frequency is the number of haplotypes harboring the tract of a specific length divided by the total number of haplotypes. *Panel* (*b*) shows the distribution of divergence between two Neanderthals captured by the ABBA-BABA (*D*) statistic in the form of (Denisovan, Altai Neanderthal, Vindija Neanderthal, Chimp) in non-overlapping 32.7 kb windows (black solid curve). The shaded gray area is defined by the lower 5% percentile of the distribution (to the left of D = 0.125). The red arrow points to the value of *D*(Denisovan, Neanderthal, Tibetan, Chimp) at the 32.7kb window within *EPAS1* identified in Huerta-Sanchez et al. ^23^. The value of *D*(Denisovan, Neanderthal, Tibetan, Chimp) in the adaptive 32.7kb region in *EPAS1* is statistically significant (*p*-value = 0.006).

High affinity to a single Denisovan genome alone, however, does not necessarily mean the introgressed segment originated from Denisovans. To examine the possibility of the beneficial haplotype in *EPAS1* instead originating from Neanderthals, we obtained the distribution of Neanderthal-Neanderthal divergence captured by computing the *D* statistic^38^ in the form of *D*(Denisovan, Altai Neanderthal, Vindija Neanderthal, Chimp) in non-overlapping 32.7kb-windows (see Supplementary Methods) to match the length of the previously identified adaptive *EPAS1* haplotype in Tibetans ^23^. This distribution, as expected, has the highest density at 1, as the two Neanderthals are more genetically related to each other (Figure 2b). If the Tibetan *EPAS1* haplotype was introduced by Neanderthals instead of Denisovans, we would expect the value of *D*(Denisovan, Neanderthal, Tibetans, Chimp) to be within the distribution because we would expect more sharing of derived alleles between Neanderthal and the Tibetan *EPAS1* haplotype. But instead, we found that the value of *D*(Denisovan, Neanderthal, Tibetans, Chimp) at *EPAS1* is significantly lower (*p*-value = 0.006), indicating that the Tibetan haplotype does not originate from a Neanderthal population, and that a Denisovan-origin for the adaptively introgressed *EPAS1* haplotype in Tibetans has the greatest support.

### The Denisovan introgression introducing the Tibetan EPAS1 haplotype occurred more than 43,000 years ago

We next sought to infer the timing of Denisovan admixture and positive selection acting on *EPAS1*. Since the introgressed haplotypes generally become fragmented over time due to recombination^39^, the distribution of introgressed tract lengths in an admixed population can be computed across the genome, and is commonly used to infer the time of admixture^40–42^. For example, simulations show that, as expected, a more recent admixture time leads to higher mean introgressed tract length^41,43^. However, some studies have suggested that selection also affects the mean introgressed tract length differently depending on whether one conditions on the present-day allele frequency^41^. Here, with simulations we confirm that the mean introgressed tract length increases with stronger positive selection when not conditioning on the current allele frequency (Figure 3, Supplementary Figure 12). This is because, under positive selection, the tract reaches high frequencies sooner while it is still long and has not been broken up by recombination. As the process of recombination then continues to break up the haplotype into more fragments, the probability that recombination results in the merger of two introgression tracts increases. In other words, the effect of selection is mediated by the allele frequency increase which elevates the probability of back-recombination between introgression fragments. Since selection acted on *EPAS1* variants, we need to account for both positive selection and archaic admixture in the modeling of the system.

**Figure 3:**
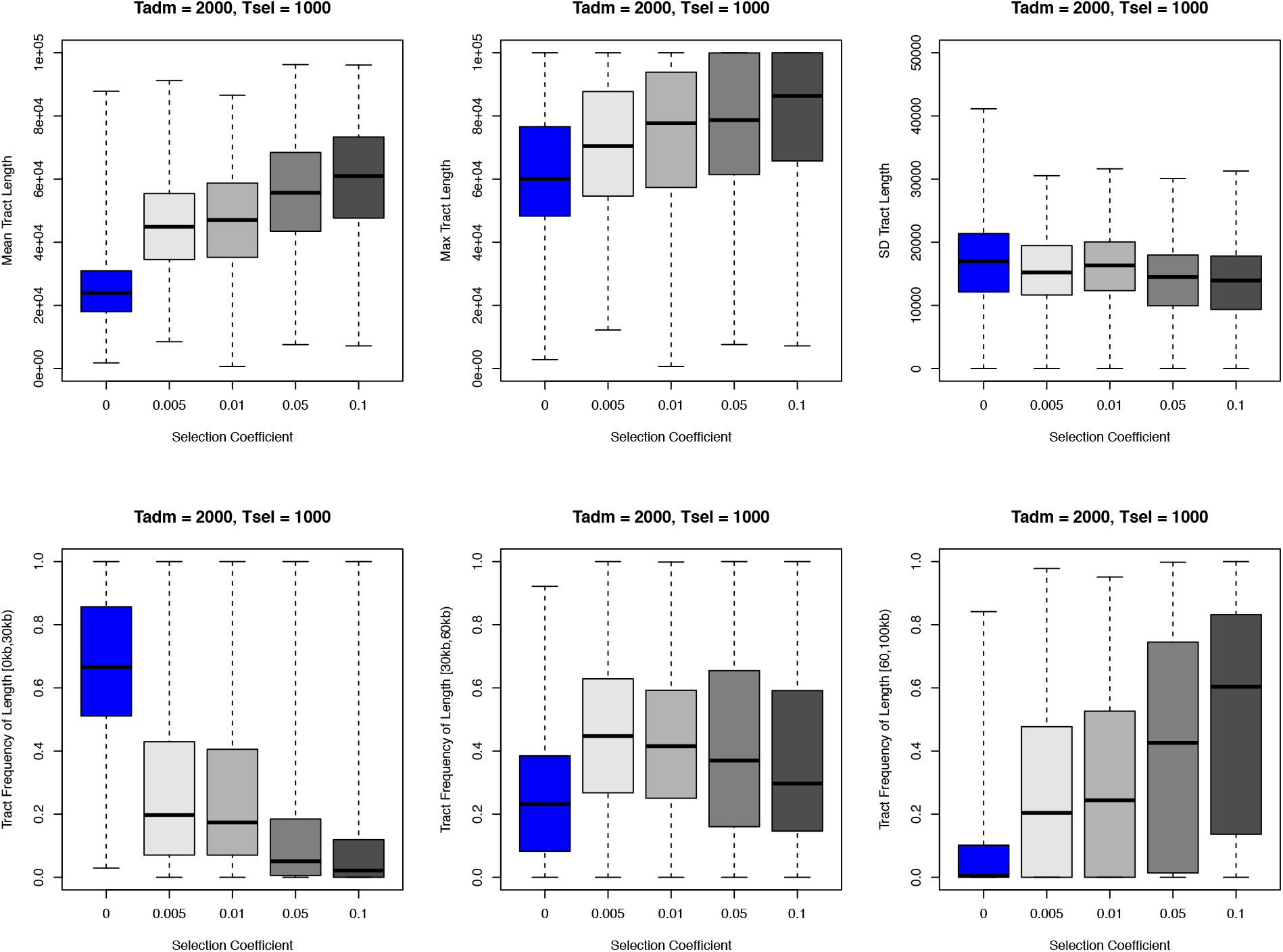
Relationship between introgressed tract length and selection coefficient, admixture time, and selection start time. We show the relationship between introgressed tract length (summarized by six statistics) and selection coefficient with simulations. In the simulations shown here, the admixture time (*T_adm_*) is fixed at 2,000 generations ago, and the selection time (*T_sel_*) starts at 1,000 generations ago (standing archaic variation). The introgressed tract lengths are tracked directly from the simulation program SLiM. Each data point in the box plot represents the statistic in all individuals from the admixed population per simulation. Each combination of evolutionary parameters (selection time, selection coefficient) was repeated 5,000 times in simulations. The demography for the simulations is the same as Supplementary Figure 7.

**Figure 4:**
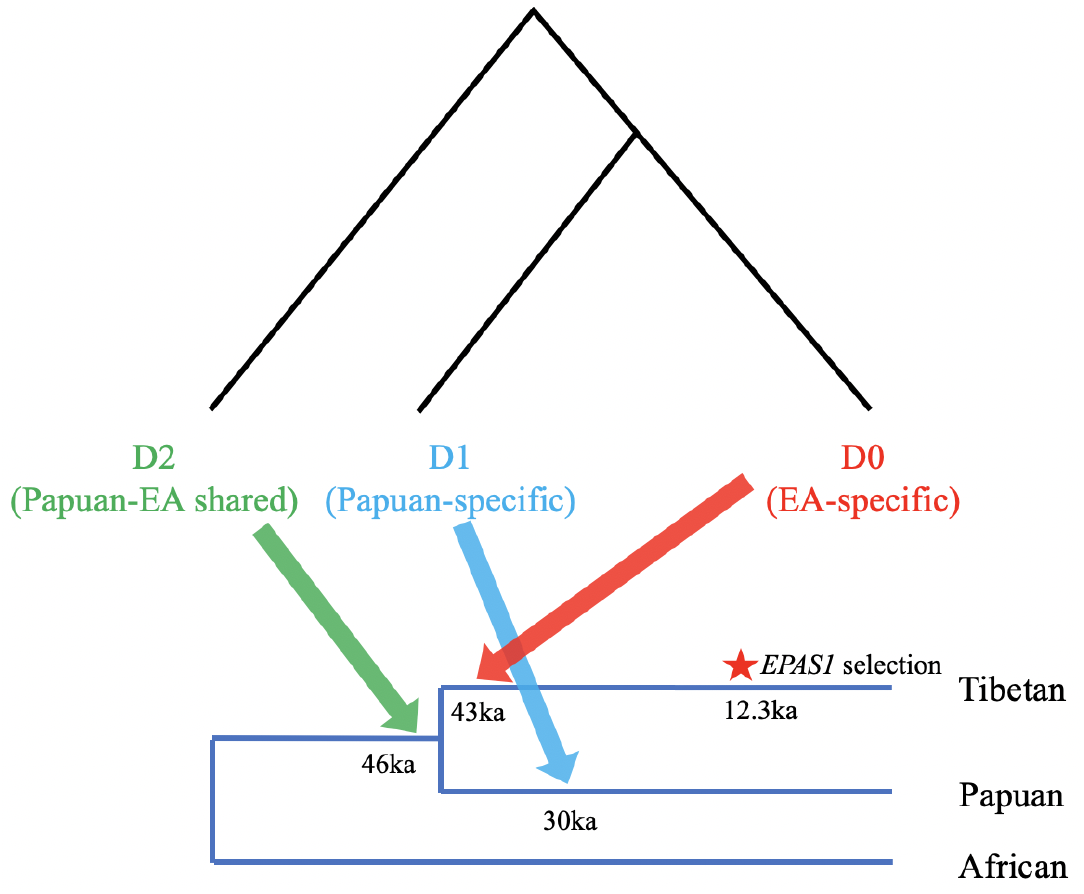
Timeline of three Denisovan introgression events in Asia, and the selection of the adaptive *EPAS1* allele. This figure is inspired by Figure 4 in Jacobs et al. 2019 that portrays three distinct Denisovan lineages (D0, D1, and D2), and their inferred introgression times. Time estimates for D1 and D2 are from Jacobs et al., and the estimate for the East-Asian specific time (D0) and the selection time in Tibetans come from this study.

We used an Approximate Bayesian Computation (ABC)-based inference framework^44–46^ and a set of summary statistics to infer three parameters: selection coefficient (*s*), the timing of selection (*T_sel_*) and admixture (*T_adm_*). We used the program SLiM3.2.0^47^ to simulate forward in time the evolution of a 100kb genomic segment representing the *EPAS1* gene under a human demographic model with Denisovan introgression (Supplementary Figure 7), using population split times estimated from Gravel *et al*.^48^ and Prüfer *et al*.^8^. At a given admixture time (*T_adm_*), a single pulse of admixture is introduced from Denisovans to the ancestral population of Tibetans at a fixed proportion of 0.1%. Subsequently, the adaptive mutation that arose in the Denisovan population remains neutral in the Tibetan population until the selection onset time (*T_sel_*). Additionally, the Tibetan population experienced a bottleneck after the split with the outgroup population, but returned to a constant size of 7,000 after 100 generations. In the simulations, the admixture time and selection coefficient were drawn from a uniform prior. The onset of selection is bounded above by the drawn admixture time, resulting in a prior that is uniform when conditioned on the admixture time.

We computed six summary statistics in both the simulations and the observed data that summarize the distribution of introgressed tract lengths. These statistics include the mean, the standard deviation, the max, and the number of tracts with length within the following three intervals: [0,30kb), [30kb,60kb), [>60kb]. Tract lengths were inferred by an HMM for each simulated chromosome and for each chromosome in the observed data at the *EPAS1* sequences from the combined dataset of 78 Tibetans (Supplementary Figure 8-9; see Methods). We first confirmed that the summary statistics chosen were informative about the parameters, especially under the demographic model we simulated. By directly tracking the introgressed segments in SLiM, we see a correlation between the statistics describing the distribution of tract length and the selection coefficient (*s*), the time of admixture (*T_adm_*), as well as the standing variation period from admixture to selection (*T_adm-_ T_sel_*) (Figure 3, Supplementary Figure 12).

We obtained a total of 400,000 simulation replicates using parameters drawn from the prior distributions and their summary statistics for the ABC inference. We used the program ABCToolBox^49^ with a rejection algorithm to retain the best-fitting 1,000 simulations for the posterior distributions. We estimated the admixture time (*T_adm_*) at 1,741 generations ago (43,525 years ago, assuming 25 years/generation; mode of posterior density; 95% credible interval [15,700 - 60,000 years ago]). The selection time estimate (*T_sel_*) is 492 generations ago (12,300 years ago; mode of posterior density; 95% credible interval [1,925 - 50,000 years ago]; Supplementary Figure 13; Table 2). Furthermore, by comparing a model of selection on standing archaic variation with selection on newly introduced archaic variants, we find more support for selection acting on standing archaic variants (Bayes factor = 2.11; Supplementary Table 1; see Methods), indicating that selection did not act immediately after introgression. The selection coefficient of the *EPAS1* haplotype (*s*) was estimated to be 0.018. Inspired by Figure 4 in Jacobs *et al*.^29^, we created a similar figure to contextualize their estimates with our estimates of Denisovan introgression events (Figure 4). Our estimates suggest that the East Asian-specific Denisovan introgression occurred at an earlier time (~43 ka) than the Papuan-specific Denisovan introgression (~30 ka).

**Table 2:** ABC estimates of *EPAS1* introgression time, selection start time, and selection strength in modern Tibetans. We used an Approximate Bayesian Computation (ABC) approach to estimate three parameters related to the evolutionary history of *EPAS1* in Tibetans, including the admixture (*T_adm_*), and positive selection start time (*T_sel_*), and the selection coefficient. We show the point estimates of parameters using the mode of the posterior distributions, and 95% credible intervals. We convert times to year units by assuming that 1 generation is 25 years.

| Parameter | Estimate | Credible Interval – 95% |
| --- | --- | --- |
| **$T_{adm}$, generations (ka)** | 1741.280 (43.53 ka) | 627.990 – 2400 |
| **$T_{sel}$, generations (ka)** | 492.055 (12.30 ka) | 77.133 – 2000.000 |
| **Selection coefficient** | 0.018 | 0.004 – 0.095 |

The bias of our ABC-based estimation method was assessed by computing the distribution of relative errors (REs, see Methods) using 1,000 randomly-sampled simulation replicates. We found that our method had highest accuracy estimating the gap period between admixture and selection start time (*T_adm_ - T_sel_*; Supplementary Figure 14). The summary statistics also show high agreement between the observed data and the retained simulations (Supplementary Figure 15). To evaluate the goodness-of-fit of our inference, we performed posterior predictive simulations (Supplementary Figure 16), which showed that the observed statistics are within the range of newly simulated summary statistics using parameters drawn from the posterior distribution.

### Archaic Introgression affects multiple genes in other biological pathways

Lastly, we investigated whether other genomic regions that were influenced by archaic introgression show signals of positive selection in Tibetans. We first asked whether the SPrime-inferred segments overlap with other high altitude adaptation candidate genes^22,28,50,51^ (Supplementary Table 2). We found a total of 11 unique regions harboring archaic segments that overlap with either a candidate gene core region, or within 200kb of the gene’s flanking region (Supplementary Table 3, Supplementary Figure 2a-b). However, most of these segments do not show signals of positive selection - for example, most SPrime-alleles except those associated with *EPAS1* and *FANCA* were not significantly differentiated between Tibetans and Han Chinese compared to their genome-wide mean *F_ST_* of 0.02^22^ (Supplementary Figure 18). This is also true for another well-known gene associated with high altitude adaptation, the *EGLN1*^22^ gene on Chromosome 1, which shows elevated *F_ST_* across the gene region, and harbors archaic alleles from Neanderthals, but shows no evidence that these archaic variants are under positive selection (low *F_ST_* on the archaic alleles). Given the evidence so far, in terms of high altitude adaptation, only the *EPAS1* gene region shows a clear adaptive introgression signal.

Next, we examined if other biological pathways received contributions from archaic introgression that facilitated positive selection. We considered all diagnostic SNPs identified from SPrime using YRI as outgroup, among which most presumably originated from Neanderthal, Denisovan, or other unknown archaic populations. The introgressed segment in the *EPAS1* region is not included in this analysis due to the concern that its exceptionally strong selection signal may dampen the weaker signals in other pathways.

Since archaic introgressed alleles are preserved in mosaic patterns on the genome, we looked for subtle signals of positive selection in subsets of a pathway by detecting enrichment of high frequency archaic alleles in genes contained in each pathway. Using the R package *signet*^52^, we identified five pathways from the National Cancer Institute / Nature Pathway Interaction Database (NCI)^53^ where the archaic alleles are enriched and potentially under positive selection (*p*-value < 0.05, Supplementary Table 4), among which two are insulin-related pathways that both contain the gene *RHOQ*. Interestingly, this gene is downstream (155kb) of *EPAS1*.

## Discussion

Previous studies have shown that archaic introgression contributed to a range of phenotypic variation in modern humans^5,54,55^, and that a number of introgressed genes were plausibly subject to positive selection^17,18^. Here, we used sequencing data of the most convincing example of adaptive introgression, *EPAS1* in Tibetans, to address a series of questions regarding the origin and the timing of Denisovan introgression in East Asia. Our work supports the two-pulse Denisovan admixture model proposed by Browning *et al*., and our analysis suggests that the beneficial haplotype of *EPAS1* in Tibetans originated from the East Asian-specific Denisovan introgression, involving a Denisovan group that is more closely related to the Altai Denisovan individual from the Denisova cave. Besides *EPAS1*, archaic introgression has left segments in various genes across the genome, and affected multiple biological pathways including hypoxia.

This work provides the first timing estimate (to our knowledge) of the East Asian-specific Denisovan introgression, which we inferred at around 43 ka, and is consistent with archaeological evidence showing Denisovan ancestry being present in modern human individuals from 34-40 ka^56,57^. Our estimate suggests that the East-Asian specific admixture event is more ancient than the Papuan-specific pulse (30 ka), and closer to the first Denisovan introgression that is shared by Asian and Oceanian lineages (45 ka, Figure 4)^29^.

The timing of human settlement in the Tibetan Plateau, including archaic hominins, is still under debate. The discovery of a partial mandible from the Middle Pleistocene (Xiahe Denisovan) in a cave located at 3,120 meters altitude in the Tibetan Plateau suggests that Denisovan-like archaic hominins may have been present at high altitude at least 160 ka^58^. Also, modern human activity has been found on the interior of the Tibetan plateau as early as 40 ka from the Nwya Devu site^59^, although long-term human settlements on the high-altitude plateau are believed to be rare at that time. Most of the archaeological evidence^60–62^ indicates that large permanent (year-round) settlements of people on the plateau started around 4,000 years ago, facilitated by the advent of agriculture. However, small groups of hunter gatherers may have lived on the plateau more distantly in the past, more than 4,000 years ago. Our estimate of the Denisovan East Asian-specific admixture time (43 ka) from tract lengths surrounding the *EPAS1* gene is larger than most estimates of when modern humans permanently settled in the Tibetan Plateau, suggesting that the admixture most likely occurred outside of this region.

Furthermore, our estimate of the onset time of positive selection on *EPAS1* (~12 ka) suggests that selection did not target the Denisovan introgressed alleles immediately after introgression, and possibly coincides with the time of permanent Tibetan settlements of populations from lowland East Asia during the Late Pleistocene or early Holocene^63^. While evidence of earlier arrivers exists (e.g. Xiahe Denisovan and the Nwya Devu site), it is unclear how long they survived in the Tibetan plateau, whether they were genetically adapted to the hypoxic environment, or if modern Tibetans are their direct descendants. Only one study has reported ancient DNA from the Himalayas (the Nepalese side) where the oldest samples date to 3.15 ka^64^. Interestingly, only the more recent samples (dated to 1.75-1.25 ka) exhibit the *EPAS1* alleles present in modern Tibetans. Future studies of ancient humans in this region should provide additional context and finer resolution to the population history on the Tibetan Plateau.

Previous studies have estimated the time of Denisovan introgression in Asia (Supplementary Table 5), but there are multiple differences between our analyses and theirs. The first relates to the underlying assumption of a single introgression event into East Asia^1,6^. Those studies used all of the surviving Denisovan segments (in Papuans or Tibetans), and it is unclear whether that estimate is an average of the two Denisovan introgression events^7,29^ into East Asians, or if the estimate is closer to one of the introgression events. By contrast, we are using the data of a single gene that clearly has been the target of selection, having the advantage that, because it is a small local region of the genome, it is highly likely that the fragments are the remnants of archaic DNA introduced by a single admixture event. The second difference is that we account for positive selection in our inference since we show that selection affects the tract length. The other estimates assume neutrality, and it is unclear whether adaptively introgressed loci could change or bias estimates from genome-wide summary statistics of introgression (e.g. the distribution of introgressed tract lengths, linkage disequilibrium decay pattern). Finally, we use an Approximate Bayesian computation framework for parameter estimation, while the estimation methods used by others^65,66^ could also lead to some differences in the inferences.

We acknowledge that we have made several assumptions and choices in our work. First, we are using the sequencing data of a single gene which is a small amount of information and that is reflected in our large credible intervals. One way to reduce uncertainty might be to use all the putative introgressed segments introduced via the East-Asian Denisovan introgression event, but doing so would require making a different set of assumptions regarding how selection is acting on each of those regions. Secondly, we have assumed a simple demographic model for Tibetans, and it is unclear how a more complex model might affect our inference. However, we did show that the bottleneck size has little effect on the admixture time (Supplementary Figure 10). Thirdly, we assumed an admixture proportion of 0.1%, and previous genome-wide estimates of Denisovan ancestry in modern-day East Asians range from 0.06%^6^ to 0.5%^14^. We chose 0.1% to ensure computational efficiency and to reduce the probability of initial loss of the beneficial mutation in the recipient population (Supplementary Figure 11). We do not expect our conclusion of selection acting on standing archaic variation to change, although using a higher admixture level will reduce the duration of the standing variation period due to a more recent estimate of the admixture time. We also do not know what the real distribution of tract lengths looks like in Tibetans, and we have inferred that using an HMM. How accurately the HMM infers the true tract lengths in Tibetans is unknown, but other methods (e.g. ArchaicSeeker 2.0^30,67^) yield similar results (Supplementary Methods & Supplementary Figure 17). Even if the HMM does not capture the true Tibetan tract lengths, by applying the HMM to both the real data and the simulated data, we hope that the same bias occurs in both, reducing the likelihood of distorting the parameter estimates.

During the last decade, we have begun to appreciate that gene flow between archaic and modern humans played a major role in shaping human evolution as well as our genetic diversity. The introduction of archaic variants evidently facilitated adaptations to local environments in multiple populations. Our results support the hypothesis that adaptive introgression mostly occurred on standing archaic variation^68,69^, and analysis of other adaptively introgressed loci will clarify whether this is the primary mode in which adaptation happens. As we continue to sequence the remains of other archaic hominins, a high-resolution picture of archaic introgression in modern humans is expected to be revealed.

## Materials and Methods

### Genomic Data from Tibetan population

For the whole genome analyses in this study, we used 38 Tibetan samples from Lu et al. 2016^30^. We phased the data with Beagle 5.0^70,71^, with the 1000 genomes worldwide populations as imputation reference. SNPs that were very rare (< 5% frequencies) or very common (>95% frequencies) were removed from the phased VCFs that are used for downstream analyses, including the SPrime analysis. For inferring the timing of admixture and selection as well as the selection strength we combined the sequences from 40 Tibetan individuals from Huerta-Sanchez et al. 2014^23^ covering a 120kb-region at the *EPAS1* gene, and the 38 Tibetan sequences at the *EPAS1* locus from Lu et al. 2016^30^.

### Genomic Data from Worldwide Population

This study utilized the following data collections as reference: (1) modern human individuals from 1000 Genomes Project^31^: Han Chinese (CHB), Yorubans (YRI), Europeans (CEU, GBR), and Peruvians (PEL), and the five South Asian populations (SAS). We also used the following archaic human genomes: Altai Neanderthal^13^, Vindija Neanderthal^8^, and Altai Denisovan^1^.

### SPrime Inference of Putative Archaic-Introgressed Genomic Segments

To infer the introgressed regions in Tibetans we used SPrime^7^. We merged the whole genome sequences of African individuals (YRI) from the 1000 Genomes Panel data to use as the reference outgroup with the VCFs of 38 Tibetans from Lu et al. 2016^30^. We ran SPrime on the combined VCF file with all 22 autosomes included to infer the putative introgressed regions in Tibetans which are each tagged by a set of sites. Alleles at these sites had maximum frequency no higher than 1% in the YRI population, as that was the threshold specified in SPrime. For each inferred segment, we further filtered out sites that were not biallelic, had low coverage depth in archaic genomes (<10), or low mapping quality score (<25). For the set of sites that passed these filters, we extracted the genotypes of the Altai Neanderthal and Altai Denisovan. For each site, we reported a “match” if the archaic genotype includes the putative introgressed allele and a “mismatch” otherwise. The match rate is calculated as the number of matches divided by the total number of sites compared (matches plus mismatches). Sites that did not pass the filters were excluded in the match rate calculation. We also applied SPrime using the CEU (individuals of European ancestry) from the 1000 Genomes Panel data as the outgroup.

To visualize the densities of match rates between the introgressed segments and the archaic genomes (in Figure 1 and Supplementary Figure 2), we utilized the function “*kde2d*” from the MASS package in R with the script from Browning et al. 2018^7^.

### Hidden-Markov Model (HMM) and the inference of introgressed tracts

Since SPrime does not identify the introgressed segments within each individual chromosome, we used a Hidden Markov Model (HMM) to call each introgressed segment in each Tibetan chromosome. We used the same HMM^13,17,37^ described in Racimo et al. (2017) and applied it to both the simulated data (under model described in Supplementary Figure 9) and the observed data. For the observed data, the HMM was used for two analyses: (1) to infer the tract lengths of the 38 Tibetans in the inferred SPrime regions whose match rate to the Denisovan was > 60% and < 40% to the Altai Neanderthal (Figure 2, Supplementary Figure 6), and (2) to infer the introgressed tract lengths at the *EPAS1* gene in 78 Tibetans (40 from Huerta-Sanchez et al. 2014 and 38 from Lu et al. 2016; see Supplementary Figure 9). In both analyses, we used 176 Yorubans (YRI) as the outgroup. We removed the non-bialleleic sites in the observed data, and formatted variants to their ancestral/derived alleles, and kept only SNPs with derived allele frequency >0 in the Tibetans. We also removed the SNPs that were private in one of the two datasets (the 40 Tibetans from Huerta-Sanchez et al. 2014 and the 38 Tibetans from Lu et al. 2016) before joining them for the HMM analysis. We plotted the inferred archaic-introgressed segments from 78 Tibetans as a heat map in the R environment. Each row represented a haplotype from Tibetan individuals, and each column represented a genomic position in *EPAS1* core region. The sites that were inferred to be archaic-introgressed were highlighted in yellow, in contrast to blue that denoted non-introgressed sites.

We further compared the results of the HMM inference on the tract length per haplotype with the inference on the 38 Tibetan individuals published by another method, ArchaicSeeker 2.0 (AS)^30,67^. ArchaicSeeker and the HMM showed high level of agreement (Supplementary Figure 17); both methods inferred a mode in intermediate-length tracts (~40kb), while HMM inferred a few more larger tracts (~80kb) and AS inferred more shorter tracts (~10kb).

### Approximate Bayesian Computation (ABC) Inference

To infer the parameters, we simulated the evolution of the *EPAS1* region under the demographic model described in the previous section (Supplementary Figure 9). We allow the admixture time (*T_adm_*) to vary between [500, 2,400] generations ago, selection time (*T_sel_*) to vary between [100, *T_adm_*) generations ago, and selection coefficient (*s*) between (0, 0.1). The simulated segment has length *L* of 100kb, which was the approximate size of the *EPAS1* region. The mutation rate *μ* was set to 1.0e-8 as estimated from Huerta-Sanchez et al^23^, and the recombination rate *r* is uniform across the segment at 2.3e-8^23^.

We applied the same HMM framework that we used for the observed data to each simulation replicate, to infer the introgressed tracts on simulated haplotypes. Given the location of HMM-inferred introgressed tract(s) in each simulation, we computed the tract length from each simulated chromosome, and recorded six summary statistics that describe the tract length distribution, including: (1) the mean (*μ*), (2) the standard deviation (*σ*), (3) the maximum length (*max*), and (4-6) the number of tracts with length within the following three intervals: [0,30kb), [30kb,60kb) and [>60kb]. The six summaries (*K*) were computed for the *EPAS1* 100-kb region in 78 Tibetan individuals. For a given parameter *θ*, the posterior probability is therefore Pr(*θ* | *K*).

We applied the program ABCToolBox^49^ to infer parameters by retaining the simulated data that best matched the observed data (as in six summary statistics). The program compares the simulated data with the observed data, and implements a rejection algorithm adjustment on retained simulations. We chose the closest 1,000 retained simulations to generate the posterior, and subsequently plotted the marginal posterior distribution of admixture time and selection start time in an R program, and obtained the mode and the 95% credible intervals (Table 2, Supplementary Figure 13, 15) from the marginal posterior distribution.

We computed the relative errors (Res, (true-estimate)/true) by randomly sampling 1000 simulations from the 400,000 set. For each of the 1,000 simulations we used ABCToolBox to infer the parameters using the rest of the simulations (a total of 399,000). We compared the difference between the inferred parameters and the true parameters (that generated the simulation replicate) for each of the 1000 randomly drawn simulations. We plotted the distribution of differences as histograms in Supplementary Figure 14. To evaluate the goodness-of-fit in our inferred scenario, we performed posterior predictive checking by randomly sampling sets of parameter values that generated 500 of the 1,000 retained simulations, and used each parameter combination to generate a single simulation, and computed the summary statistics. We plotted the relationship of pairwise summary statistics between the observed data, the sampled 500 retained simulation, and the generated 500 simulation replicates under the 500 parameter combinations, and show that the summary statistics from the observed data are within the range of both retained posterior and the newly simulated data (Supplementary Figure 16).

Lastly, we performed model selection using ABCToolBox (Supplementary Table 1). We reran the inference of ABC using the simulations generated previously under the model of selection on standing archaic variation (M1), together with an additional 400,000 simulations generated under a model of immediate selection on archaic variation (M2), where the only difference is that in M2, selection acts immediately after introgression in the Tibetan population. The Bayes factor^72^, or the ratio of posterior probabilities of the two competing models with equal prior is 2.11, suggesting more support for M1 than M2.

### Biological Pathway Network Analysis

We looked for subtle signals that a subset of genes within a pathway network may be selected, by searching for enrichment of archaic alleles in gene networks in biological pathway databases and finding the highest scoring subnetwork^52^. First, we searched for overlaps between the putatively introgressed segments inferred by SPrime and protein-coding genes in modern humans using the ENSEMBL database^73,74^, which resulted in 3,292 genes by combining the searches using either the YRI or CEU as outgroup populations. At the overlapping SPrime diagnostic SNPs, we calculated their frequencies in the Tibetan population. We used the maximum archaic allele frequency in each gene as input score for the enrichment test. Alternatively, if no archaic allele was found in a gene, the gene received a score of 0. We then computed the subnetwork scoring using the National Cancer Institute/Nature Pathway Interaction Database (NCI)^53^ as reference for signaling and metabolic pathways, using the HSS algorithm provided under R package *signet*^52^. The final score of each subnetwork was normalized using the mean and standard deviation of 10,000 simulated random networks of the same size. We reported only the significant subnetworks with *p*-values less than 0.05 as candidate biological pathways that undergo positive selection because of archaic allele enrichment (Supplementary Table 4).

## Data Availability Statement

All scripts necessary to reproduce the ABC and simulation results from this work can be found on https://github.com/xzhang-popgen/EPAS1Project. The use of 38 Tibetan whole genomes by this work is permitted by The Ministry of Science and Technology of the People’s Republic of China (Permission No. 2020BAT0143). The *EPAS1* sequences of 40 Tibetans are publicly available at Sequence Read Archive under accession number SRP041218.

## Acknowledgement

This study was supported by an NSF grant # 1557151 and NIH grant 1R35GM128946-01 (to E.H.S.). X.Z. was partially supported by NIH grant R35GM119856 to Kirk Lohmueller. S.X. gratefully acknowledges the support of the National Natural Science Foundation of China (NSFC) grant (31525014, 91731303, 31771388, 31961130380, and 32041008), the Shanghai Municipal Science and Technology Major Project (2017SHZDZX01), and the UK Royal Society-Newton Advanced Fellowship (NAF\R1\191094). The authors thank the Huerta-Sanchez lab at Brown University and the Lohmueller lab at UCLA for helpful discussions during the development of this work.

