## Supplementary Material for "The history and evolution of the Denisovan-*EPAS1* haplotype in Tibetans"

**Supplementary Methods, Figures and Tables**

ABBA-BABA ($D$ statistics) distribution

To reject the hypothesis that the Tibetan EPAS1-haplotype was from Neanderthals, we calculated the distribution of $D$ statistics of the form $D$(Denisovan, Altai Neanderthal, Vindija Neanderthal, Human-Chimp Ancestor) using non-overlapping 32.7-kb windows across the genome (excluding telomeres and centromeres). We chose 32.7kb because this is the length of the haplotype identified in Huerta-Sanchez et al. (2014). We removed archaic sites that had low genotype quality score (<40) and low mapping quality (<30). Within each window, we used two methods of computing the $D$ statistic (Durand et al. 2011[1]). First, we randomly sampled an allele from each of the archaic human haplotypes at each side to find sites that matched the ABBA or BABA allele pattern. Second, we used the equation that incorporates allele frequencies to estimate the $D$-statistic for windows with few ABBA and BABA sites. Using this method, we calculated the allele frequency of the derived allele for each individual (1, 0.5, or 0 for 2, 1 or 0 copies of the derived allele respectively) and used those frequencies to make the calculations. The values of $D$ statistic obtained across the genome were shown as a density distribution curve in Figure 2b.

Similarly, we computed the value of $D$(Denisovan, Neanderthal, Tibetan, Chimp) at the 32.7kb region in *EPAS1* (chr2:46,567,916:46,600,661, hg19) using the allele frequencies of derived alleles in each population, with Tibetan population being a joined dataset of 78 individuals (40 from Huerta-Sanchez et al. 2014 and 38 from Lu et al. 2016). Using Altai or Vindija individual as Neanderthal yielded the same $D$ statistic value, and we show it as a red solid arrow in Figure 2b. We additionally highlight the $D$(Denisovan, Altai Neanderthal, Vindija Neanderthal, Human-Chimp Ancestor) at this window within *EPAS1* as a blue solid line (Figure 2b). We computed the $p$-value of the $D$(Denisovan, Neanderthals, Tibetan, Chimp) at *EPAS1* with Tibetans in the tree using the genome-wide distribution.

Forward Simulations in program SLiM

We used SLiM 3.2.0[2] to perform all simulations in this study. To reduce the computational burden of simulations, we rescaled the simulation parameters in this study by a scaling factor $C$, where $C$=10 in this study. The principle of scaling the parameter followed: population size = $N/C$, times = $t/C$, selection coefficients = $s\times C$, mutation rate = $\mu\times C$, and recombination rate = $r\times C$. The total length of simulated genomic sequence remained the same. Throughout the paper, we describe the simulation parameters as original parameters before scaling.

We simulated under the three-population demographic model described in Supplementary Figure 7. After a burn-in period of $10\times N$ generations (100,000 generations in this study) of a single population representing the ancestral population of modern and archaic humans. At 16,000 generations ago, the population splits into an archaic population (Denisovans) and the ancestral modern human population. After that, at 4,000 generations ago, ancestral modern human splits into the ancestral Tibetan population and African population. At this split the ancestral Tibetan population reduces to a size of 120 individuals. The bottleneck lasts for 100 generations, and then the population recovers to a size of 7,000. Subsequently, at a time point drawn from a uniform distribution between 500 generations ago to 2,400 generations ago a single pulse of gene flow occurred from Denisovans into ancestral Tibetans. The admixture proportion was set to 0.1%.

We introduced 1 adaptive mutation in the middle of the simulated segment at 15,000 generations ago to all haplotypes in the archaic population to ensure fixation in the archaic population before introgression. The selection coefficient (*s*) of the adaptive mutation varied between 0 and 0.02 with step size of 0.0002. Right after admixture, the previously adaptive mutation (in the archaic population) was set to be neutral in Tibetans until the specified onset time of selection. For each admixture time, a randomly chosen selection start time was simulated between right after admixture (up to 2,400 generations/72,000 years ago) and as soon as 100 generations ago (3,000 years ago), at average step size of 100 generations. At the time of selection, the selection coefficient of the adaptive mutation in Tibetans resumed to its original value as when introduced in Denisovans, and remained constant until the end of the simulation. We only kept the simulations where the adaptive mutation was not lost in the recipient population by the end of each simulation (adaptive mutation frequency > 0 in Tibetans). We sampled 2, 176, and 156 haplotypes from the Denisovan, African and Tibetan populations respectively, matching the sample size in the empirical dataset of Denisovans, unadmixed Africans (YRI), and Tibetans.

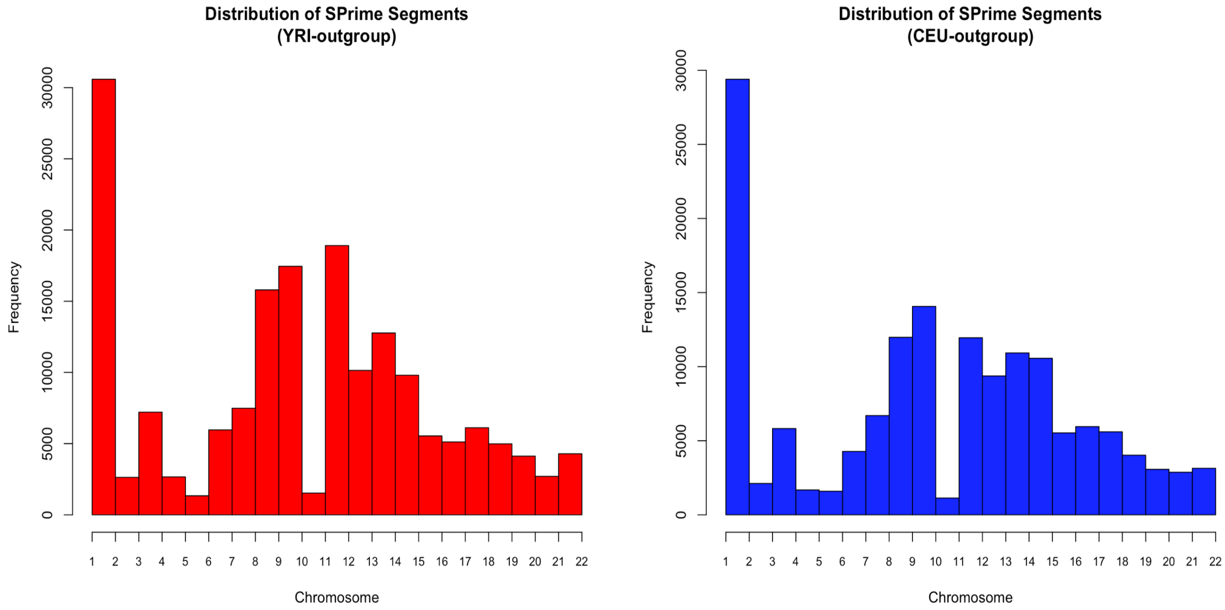


**Supplementary Figure 1: Chromosomal distribution of putatively archaic-introgressed segments inferred using YRI and CEU as outgroup.** *This figure shows the distribution of putatively introgressed segments on the 22 autosomes, inferred by SPrime using YRI and CEU as outgroup respectively. The X-axis denotes the chromosome number. The Y-axis is the number of segments in the corresponding chromosome.*

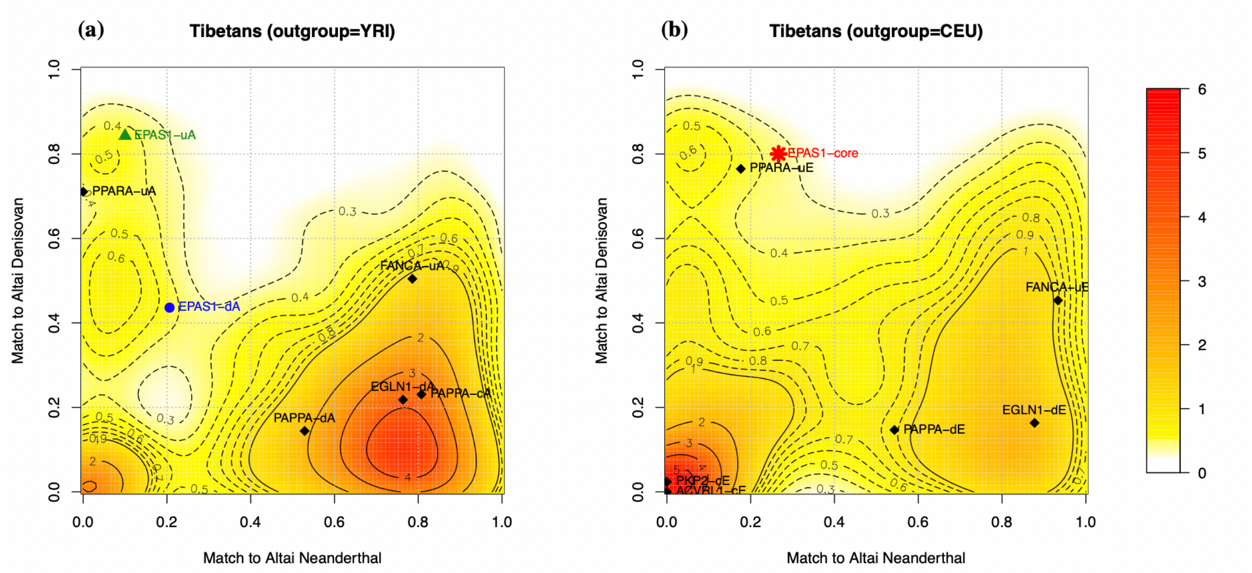


**Supplementary Figure 2: Match rate distribution of introgressed segments in Tibetans.** *This figure shows the density distribution of match rate to archaic individuals in putatively archaic introgressed segments in Tibetan population (n=38), inferred by SPrime using Africans as unadmixed outgroup (YRI, panel a) or using Europeans as the outgroup (CEU, panel b). The match rate is defined as the proportion of alleles at the SPrime diagnostic SNPs in a putatively*

*introgressed segment that are present in the genome of archaic individuals at those positions* [3]. *The color range denotes the density of the contours, with red indicating high density and yellow indicating low density. The introgressed segment detected within the EPAS1 gene is highlighted as a star symbol. The introgressed segments detected at or within 200kb range of other high altitude adaptation candidate gene (Supplementary Table 3) are highlighted as points. After each gene name, "-u/c/d" indicates the location of the segments being in the gene upstream, core, or downstream region respectively. "A" or "E" indicates whether the segment was inferred using Africans (YRI) or Europeans (CEU) as reference.*

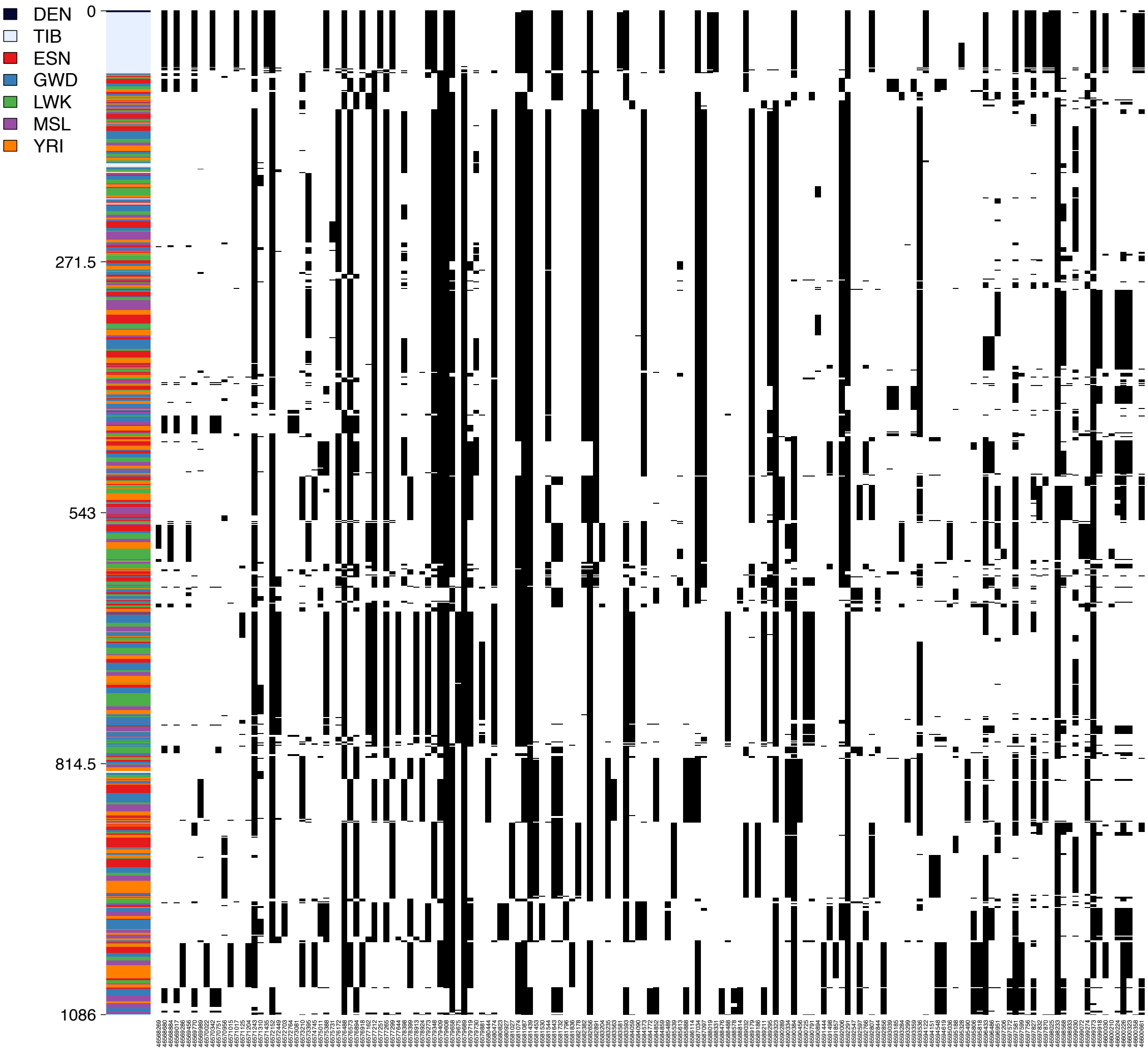

**Supplementary Figure 3. Denisovan-introgressed haplotype at *EPAS1*.** *We show the haplotype patterns of the introgressed segment in the core region of EPAS1 in Denisovan, Tibetans, and African populations in 1000 Genomes Project using haplostrips program[4]. Each column corresponds to a SNP with black cells representing the presence of derived alleles. Each row is a phased haplotype. The colored panel on the left shows the population for haplotypes. Unphased Denisovan genotypes are shown as the top two rows (black).*

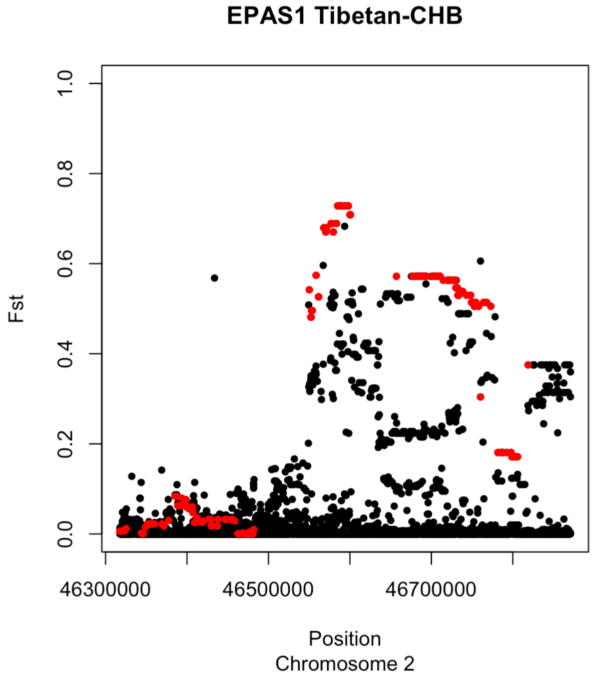


**Supplementary Figure 4: $F_{ST}$ between Tibetans and Han Chinese within 200kb of *EPAS1* region.** *We show the $F_{ST}$ values of all SNPs at or within 200kb region of EPAS1 in black points. The x-axis shows the genomic position of the SNPs (hg19), and the y-axis shows the $F_{ST}$ value. The red points highlight the diagnostic SNPs that are tagged in this region detected by SPrime (See Table 1). $F_{ST}$ values are calculated between Tibetan and Han Chinese (CHB). The introgressed variants within the EPAS1 region show the highest allele frequency differentiation, indicating positive selection.*

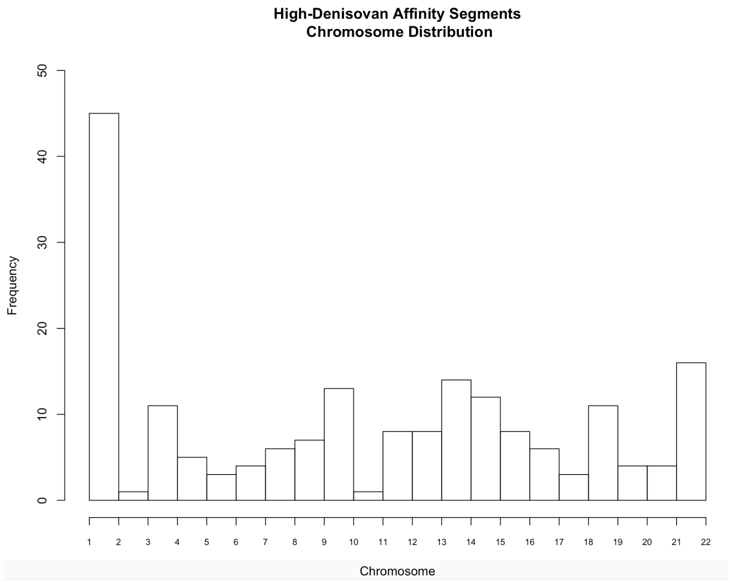


**Supplementary Figure 5. "High Denisovan Affinity" introgressed segment distribution on 22 autosomes.** *We show the chromosomal distribution of SPrime segments that have match rate less than 40% to Altai Neanderthal, and larger than 60% to Altai Denisovan, which represent the segments introduced by the East Asian-specific Denisovan introgression.*

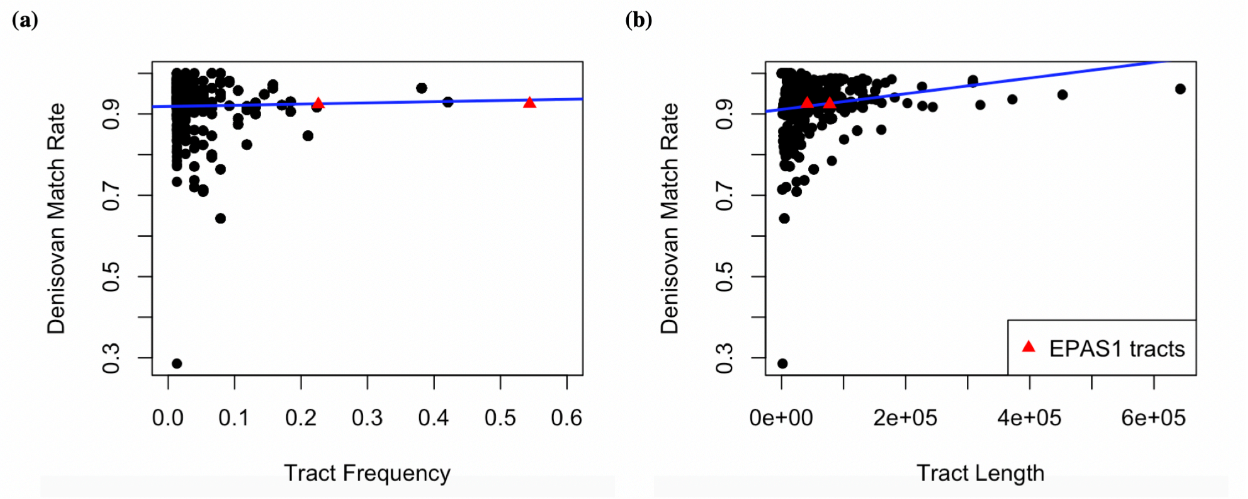


**Supplementary Figure 6. Introgressed tract frequency, length and Denisovan match rate relationship.** *This figure shows match rates distribution of Denisovan-introgressed segments, with respect to the frequency of each segment in Tibetan population (panel a), and the length of these tracts (panel b). All segments are detected by HMM from 38 Tibetan whole genomes. The black points represent introgressed tracts inferred by HMM, summarized by tract length. The input genotypes for HMM inference include derived alleles within the range of the first and last diagnostic SNPs in each unique SPrime segment. The red triangle points represent the introgressed tracts at EPAS1, including a long and a short segment (80kb and 40kb respectively, see Supplementary Figure 9). The tract length is computed by the distance between the first and last position of a continuous introgressed tract inferred from each haplotype. The tract frequency is the number of haplotypes harboring the tract of a specific length divided by the total number of haplotypes. The match rate is defined as the proportion of SNPs in a segment that is present in the Altai Denisovan genome at respective positions.*

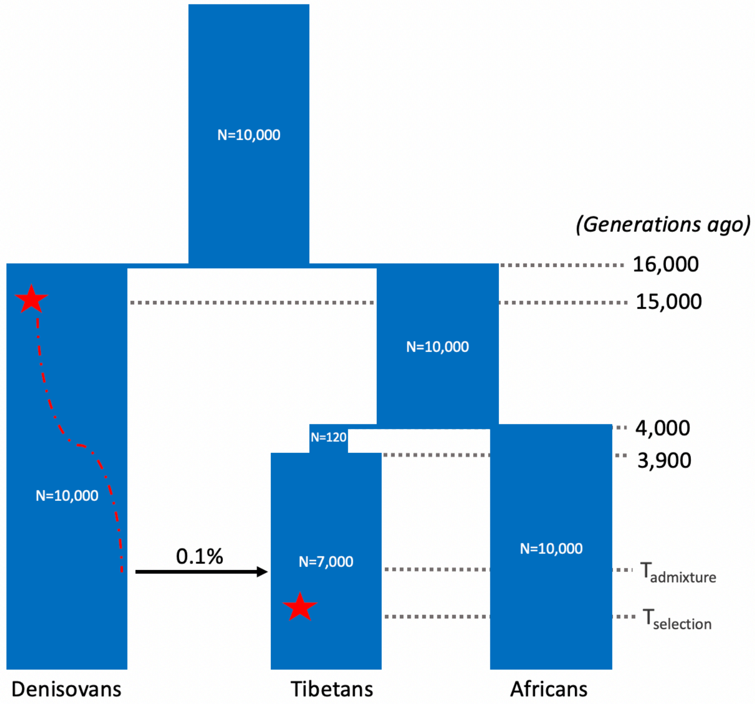


| Parameter | Value Range |
|---|---|
| Admixture time ($T_{admixture}$) | [500, 2400] |
| Selection start time ($T_{selection}$) | [100, $T_{admixture}$) |
| Selection Strength | [0, 0.1] |

**Supplementary Figure 7: ABC Simulation demography model and parameters.** *Going forward in time, after a burn-in period of 10*N (100k) generations, the ancestral population split into two subpopulations, the archaic (Denisovans) and ancestral modern human populations, at 16k generations ago (g.o). At 15k g.o, an adaptive mutation occurred in Denisovans and became fixed before introgression. The ancestral modern human split into Non-Africans (Tibetans) and Africans at 4k g.o. After 100 generations of a bottleneck, the Tibetan population resumed population size, and later received a pulse of introgression such that 0.1% of the ancestry in Tibetans came from Denisovans. The adaptive mutation on EPAS1 went through a neutral period before being selected again in Tibetans.*

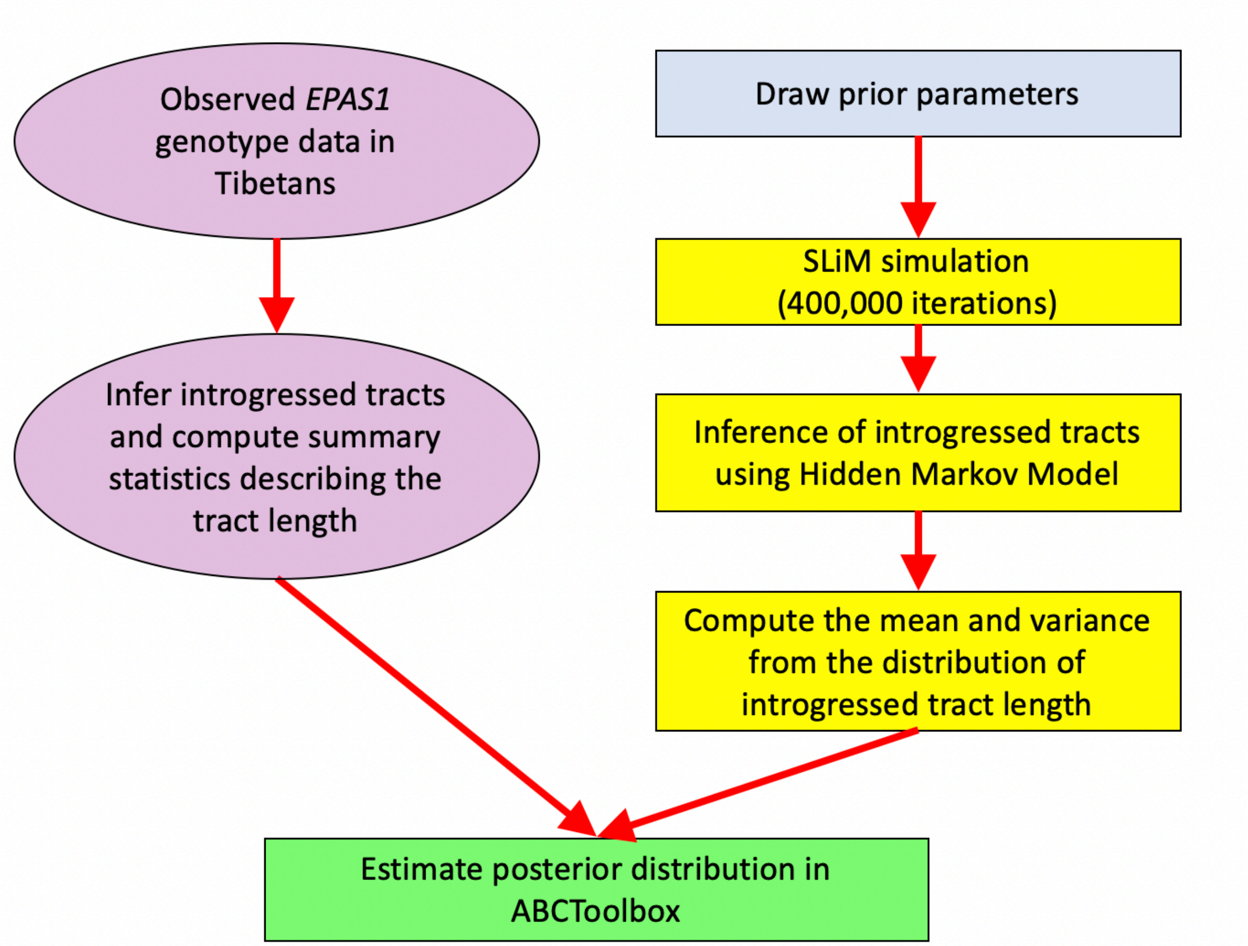


**Supplementary Figure 8: Flowchart illustration of the ABC inference framework**

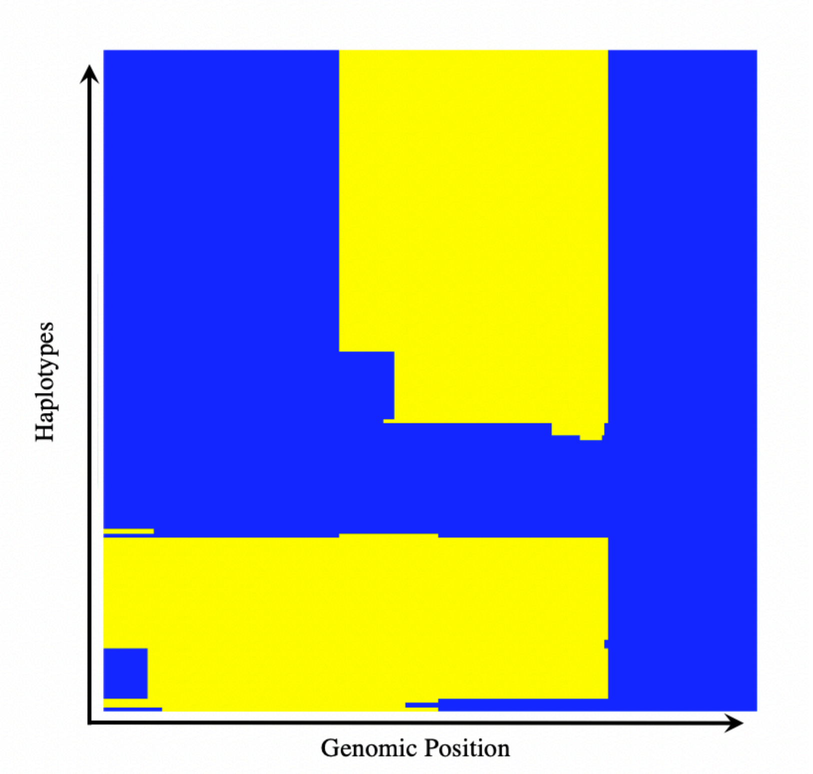


**Supplementary Figure 9: HMM-inferred introgressed tracts in the haplotypes of 78 modern Tibetans at *EPAS1* region.** *This figure shows the distribution of introgressed tracts in haplotypes of the combined set of 78 Tibetans at EPAS1, inferred by HMM. The introgressed tracts are highlighted in yellow in respect to the EPAS1 region as blue background. The x-axis shows the genomic position of the tracts, and each horizontal line along the y-axis shows a Tibetan haplotype (156 total).*

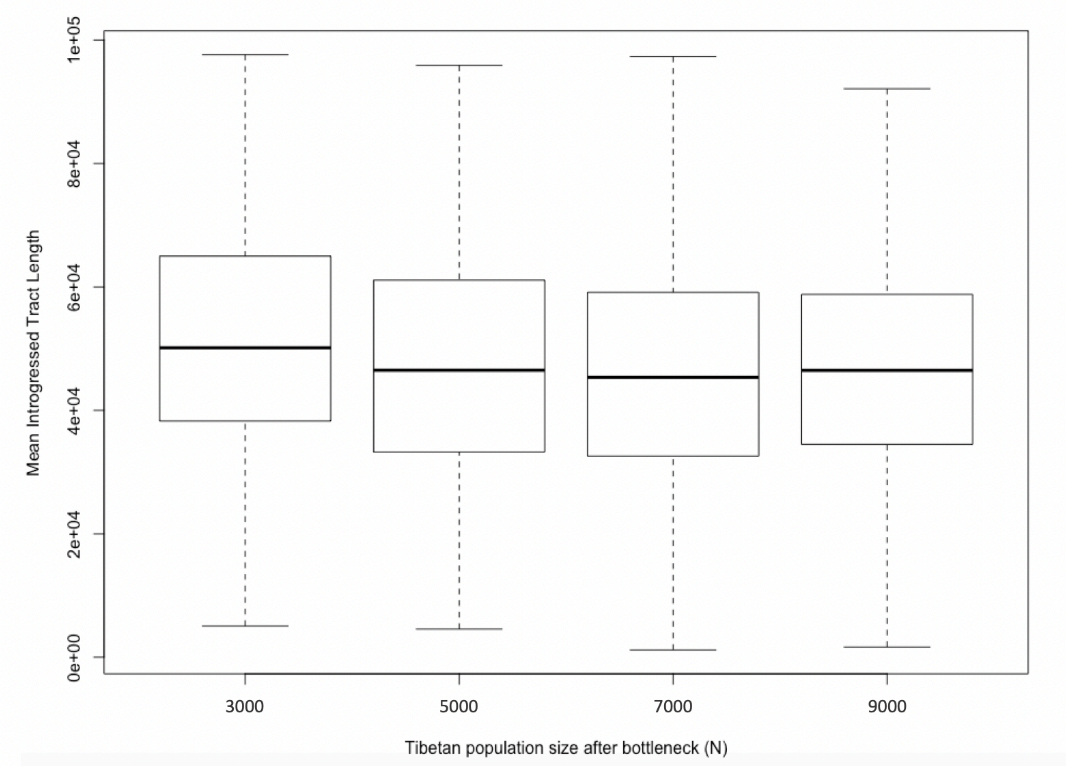


**Supplementary Figure 10: Relationship between Tibetan population size after the bottleneck and introgressed tract length inference.** *This figure shows the relationship between the mean introgressed tract length and the resumed Tibetan population size after a bottleneck in the simulations (N = 3,000, 5,000, 7,000, and 9,000 respectively). The effective population size in Tibetans after the out-of-Africa bottleneck (N=120) shows little correlation with the mean introgressed tract length. We used N=7000 as the population size by the time Tibetans received introgression from Denisovans.*

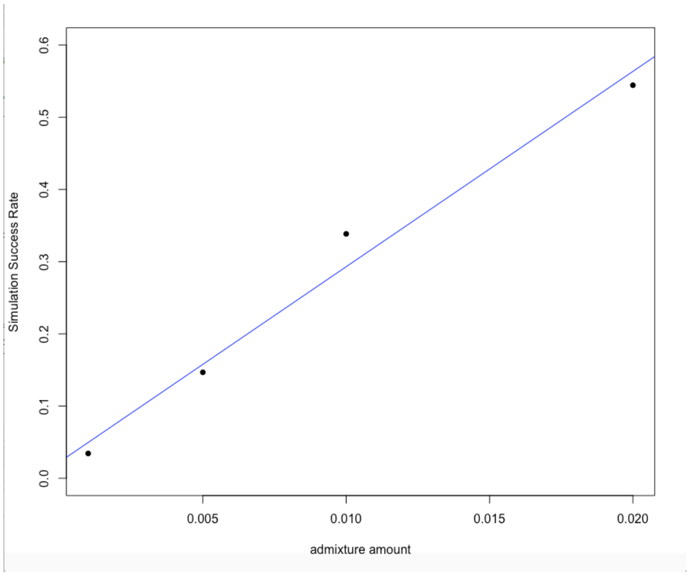


**Supplementary Figure 11: Relationship between successful simulation rate and the admixture proportion.** *This figure shows the relationship between the admixture amount from Denisovans into Tibetans in the simulations and the simulation success rate. Simulation success rate is defined as the proportion of simulation replicate with the introgressed adaptive mutation still segregating in the Tibetans by the end of the simulation, with respect to the total number of simulation attempted (N=1,000 in our study). We chose a fixed rate of 0.1% admixture proportion to leverage the simulation efficiency and empirical estimate of genome-wide Denisovan ancestry in East Asians (0.06%-0.5%).*

**(a) Selection on Standing Archaic Variation**

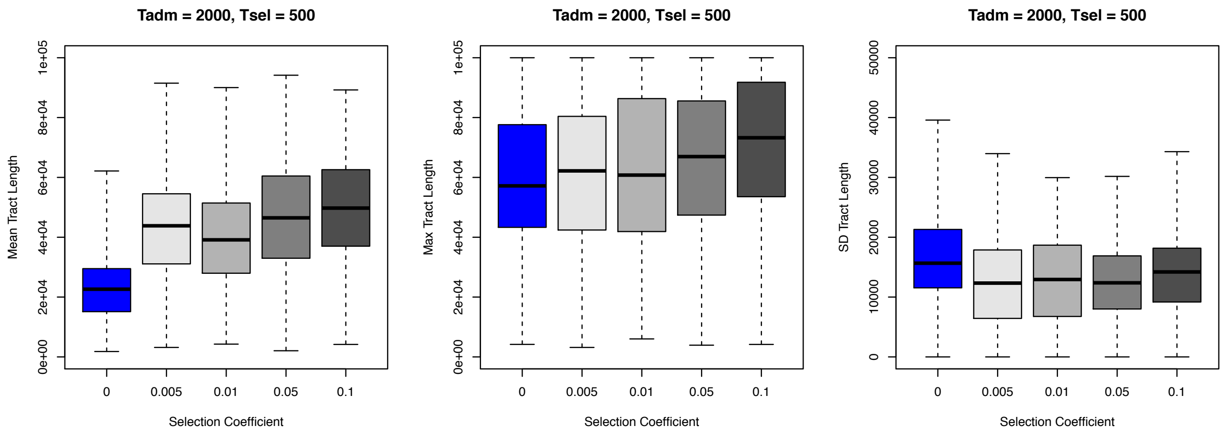


**(b) Adaptive Introgression (no neutral period)**

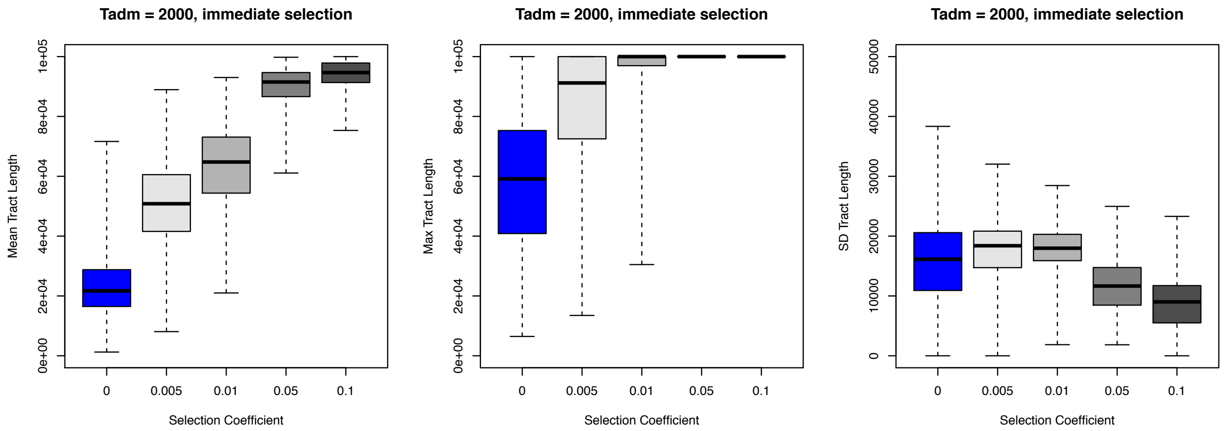


**Supplementary Figure 12: Relationship between introgressed tract length and selection coefficient, admixture time, and selection start time.** *We show the relationship between introgressed tract length (summarized by six statistics) and the strength of selection in simulations used for ABC inference. In the simulations, the admixture time (Tadm) is fixed at 2,000 generations ago, and the selection time started either at 500 generations ago (panel a; selection on standing archaic variation), or immediately after the introgression (panel b; adaptive introgression). The introgressed tract lengths are tracked directly from the simulation program SLiM. Each data point in the box plot represents the statistic in all individuals from the admixed population per simulation. Each combination of evolutionary parameters (selection time, selection coefficient) was repeated 5,000 times in simulations. The rest of the demography for the simulations is the same as Supplementary Figure 7.*

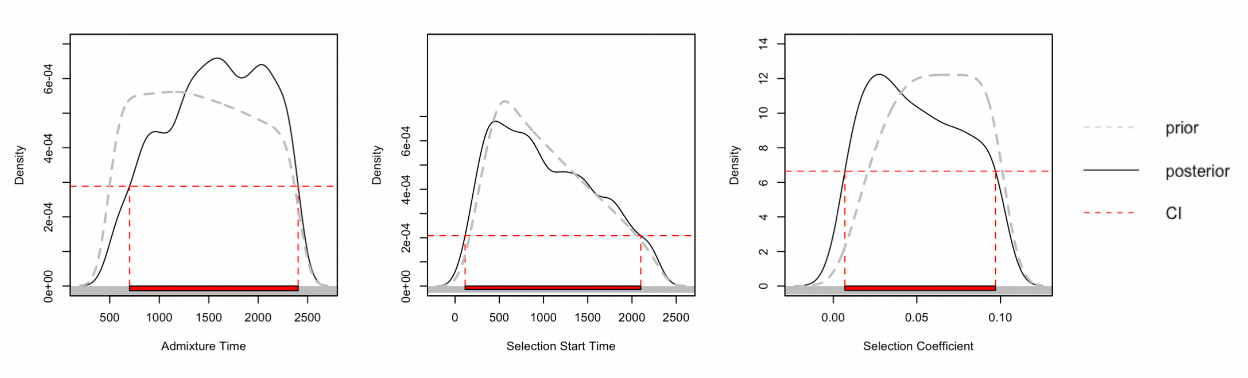


**Supplementary Figure 13: ABC Parameter Estimates.** *From left to right, the panels show the ABC parameter estimates for the admixture time, selection start time, and selection coefficient, inferred from simulations using 0.1% admixture proportion. In each panel, the gray dotted line shows the density distribution of parameters in the simulations (prior), the black solid line shows the density distribution of 1,000 retained simulations from ABC using rejection algorithm (posterior), and red dotted line encloses the 95% credible interval range. For the admixture and selection start time, the unit for the values is "generations ago", where one generation is assumed to be 25 years.*

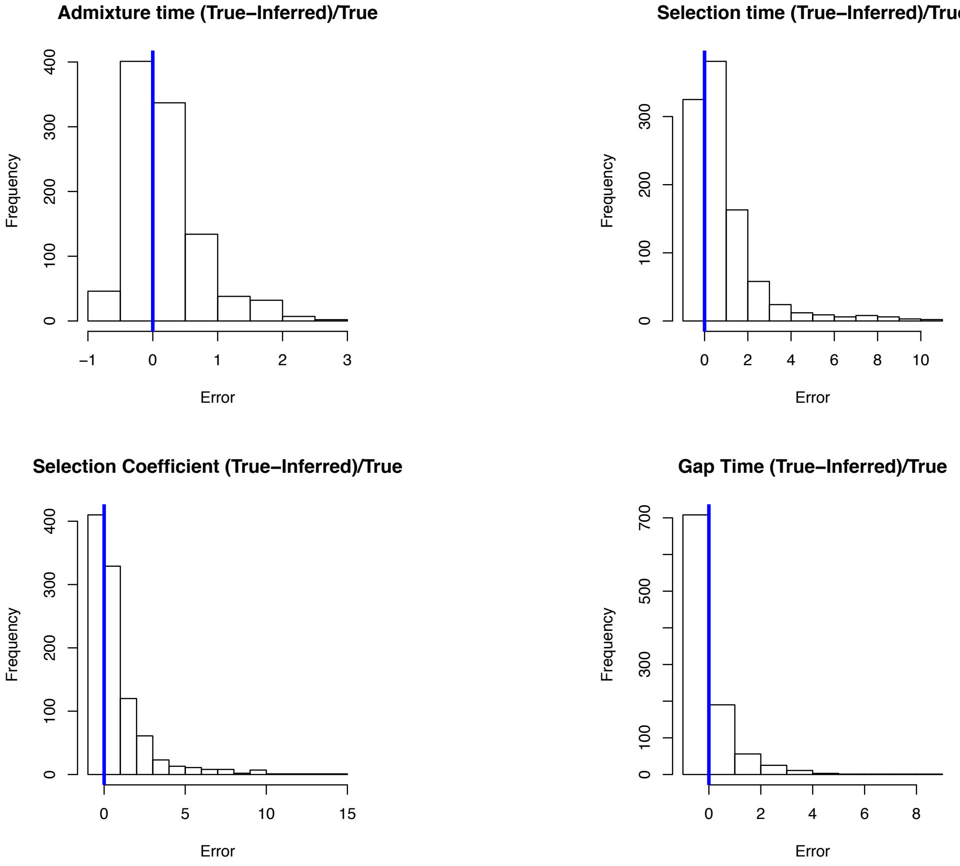


**Supplementary Figure 14: ABC Parameter estimate error distribution from 1,000 randomly chosen simulations using 0.1% admixture amount.** *In this figure, we vetted the accuracy of the ABC approach by randomly drawing 1,000 simulations from the total of 400,000 simulations (0.1% admixture), and inferred the parameters (admixture time, selection time, selection coefficient) in ABCToolBox using the rest 399,000 simulations. We compared the difference between the inferred parameters and the true parameters in the 1,000 set, and computed "relative errors" (REs) for each summary statistics where RE = (true parameter – inferred parameter)/true parameter. The above figures show the distribution of the errors in admixture time, selection time, selection coefficient, and the difference between admixture and selection time (Gap Time). We show small biases in estimating the parameters, with particularly high accuracy in detecting the gap time between admixture and selection.*

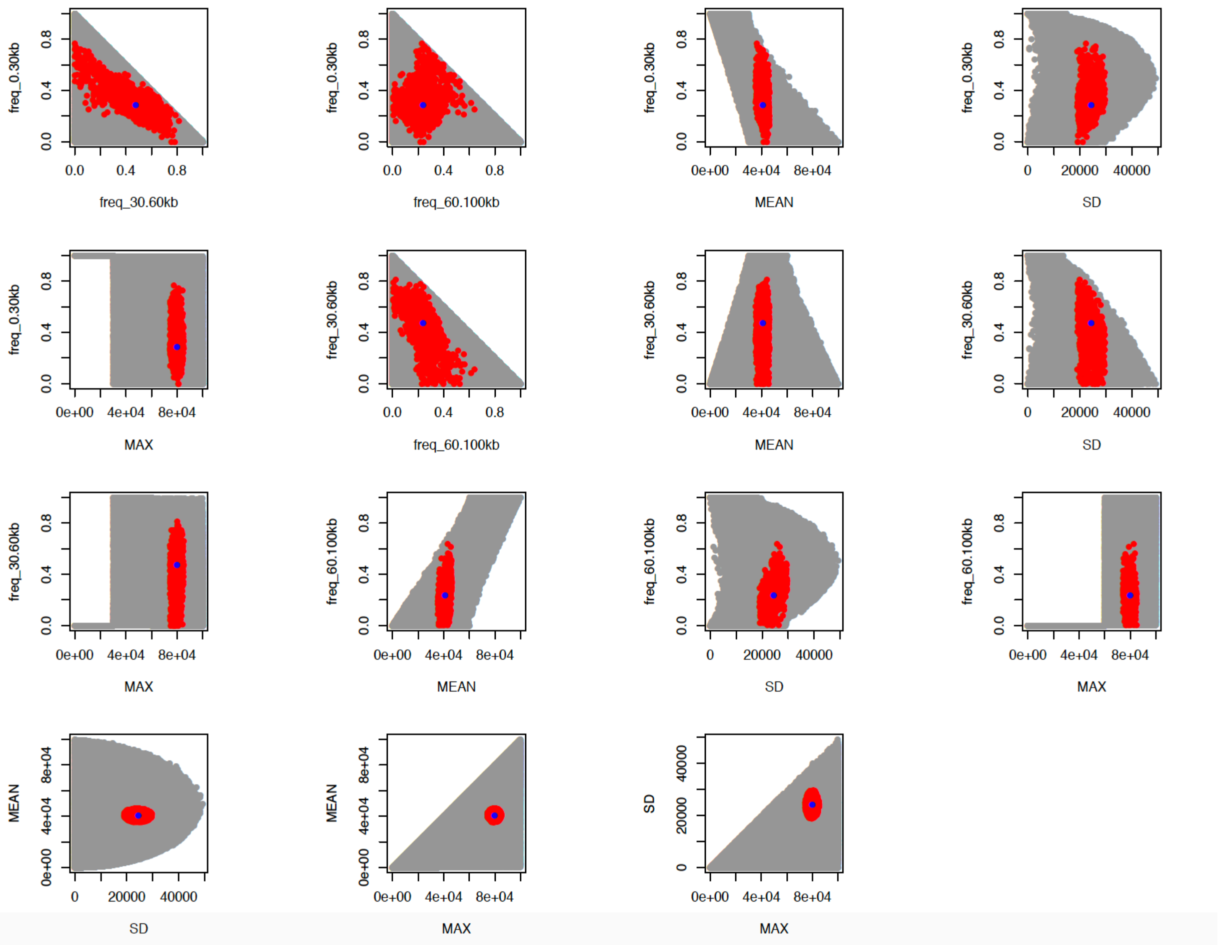


**Supplementary Figure 15: The pairwise distribution of summary statistics used in ABC inference from observation, simulations, and retained best simulations.** *This figure shows the distribution of pairwise summary statistics between the simulated data (gray points), retained 1,000 best fitting simulations (red points), and the observed data from 78 Tibetans (blue points). The retained statistics and the observed data show high agreement.*

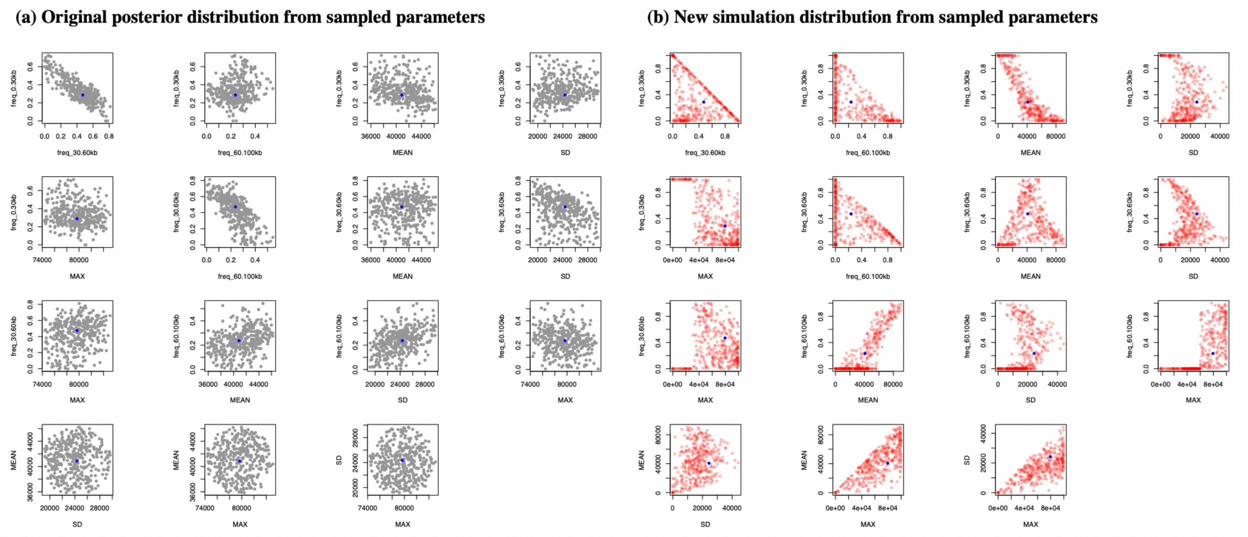


**Supplementary Figure 16: Posterior predictive checking of 500 randomly sampled sets of parameters from retained posterior.** *In this figure, we performed posterior predictive checking of the ABC inference by randomly sampling 500 set of simulation parameters from the retained posterior (n=1,000). For each set of parameters, we obtained 1 new simulation using these parameters, and computed the summary statistics in ABC. We compared the original distribution of pairwise summary statistics from the sampled posterior (left panel, posterior in gray points) with regards to the observed data (blue points), and the distribution of pairwise statistics obtained from the new simulations (right panel, new simulations in red points) with regards to the observed data. We show that parameters sampled from the posterior are capable of reproducing the range of summary statistics seen in the posterior.*

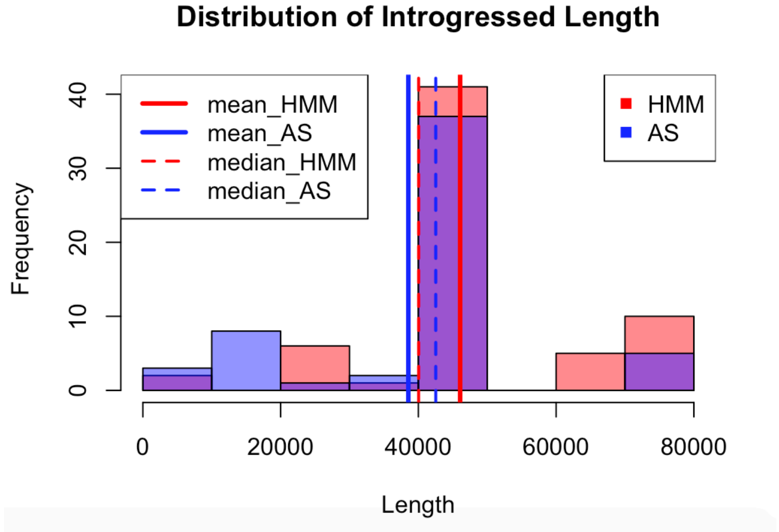


**Supplementary Figure 17: Distribution of introgressed tract length in *EPAS1* region inferred by HMM and ArchaicSeeker 2.0[5].** *In this figure, we compared the distribution of introgressed tract length inferred from 38 Tibetans at haplotype level within the EPAS1 region, inferred from HMM (red) and ArchaicSeeker 2.0 ("AS", blue) programs. The solid and dashed lines highlight the mean and median of tract length respectively. We show high level of agreement between the two methods for length inference.*

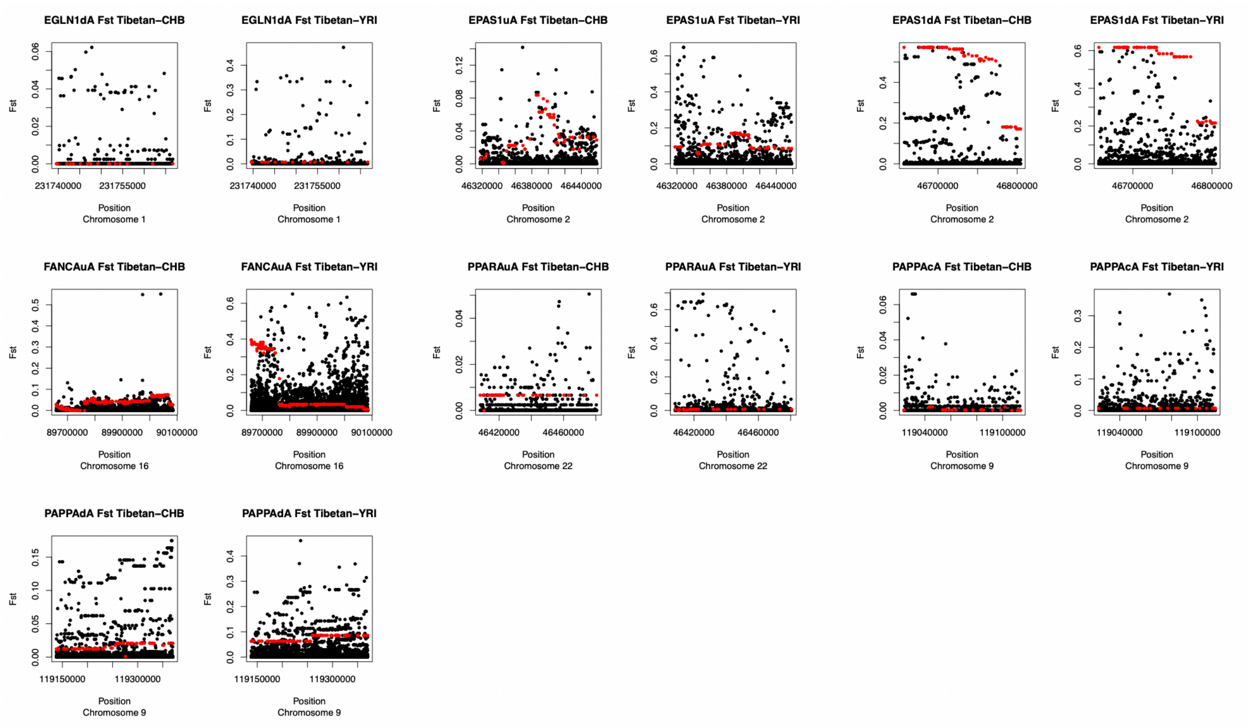


**Supplementary Figure 18: $F_{ST}$ between Tibetans, Han Chinese and Yorubans on segments overlapping between high altitude adaptation candidate genes and SPrime-inferred segments.** *The above figure show the $F_{ST}$ values of SNP variants (black points) within SPrime-inferred segments that are at or near high altitude adaptation candidate genes, with introgressed archaic variants highlighted as red points (archaic variants = diagnostic SNPs inferred from SPrime). After each gene name, "u/c/d" indicates the location of the segments being in the gene upstream, core, or downstream region respectively. "A" indicates that these segments were inferred using Africans (YRI) as the outgroup.*

**Supplementary Table 1: Model choice estimates**

| M1 Marginal Density | M2 Marginal Density | M1 Posterior Probability | M2 Posterior Probability | Bayes Factor M1/M2 | Chosen Model |
|---|---|---|---|---|---|
| 2.00E-20 | 9.51E-21 | 0.678135 | 0.321865 | 2.10689 | 1 |

*This table shows the model selection bias between two competing models for the selection of Denisovan EPAS1 allele in Tibetan populations: "Selection on standing archaic variation" model (M1) where a gapped time between introgression and selection is observed, and "Immediate selection on archaic variation" model (M2) where the positive selection is continuous before and after the introgression. We obtained equal number of simulation replicates between the two models (n=400,000), and inferred posterior probabilities using*

*ABCToolBox. The Bayes Factor, which measures the ratio between posterior probabilities of two models, shows a strong favor for M1 over M2.*

**Supplementary Table 2: High Altitude Adaptation-Associated Genes (Hg19/GRCh37 Coordinates)**

| Chromosome | Start Position (hg19) | End Position (hg19) | Gene Name |
|---|---|---|---|
| 1 | 155034154 | 155496154 | *PKLR* |
| 1 | 158373495 | 158863506 | *SPTA1* |
| 1 | 176225306 | 177018970 | *PAPPA2* |
| 1 | 231292497 | 231767790 | *EGLN1/DISC1* |
| 2 | 46317540 | 46820842 | *EPAS1* |
| 2 | 108656650 | 109211270 | *SULT1C3* |
| 2 | 203896163 | 204503892 | *CYP20A1* |
| 5 | 58057865 | 59990925 | *PDE4D* |
| 6 | 25860489 | 26322489 | *HFE* |
| 6 | 71170478 | 71778716 | *SMAP1* |
| 9 | 118709070 | 119371600 | *PAPPA* |
| 10 | 89416194 | 89935532 | *PTEN* |
| 10 | 104362789 | 104824789 | *CYP17A1* |
| 11 | 5038034 | 5500034 | *HBB/HBE* |
| 12 | 32736679 | 33256780 | *PKP2* |
| 12 | 33321347 | 33799754 | *SYT10* |
| 12 | 52078173 | 52540173 | *ACVRL1* |
| 12 | 120220647 | 120739299 | *CCDC64* |
| 15 | 48192879 | 48654879 | *SLC24A5* |
| 16 | 4311533 | 4773533 | *HMOX2* |
| 16 | 89596958 | 90090065 | *FANCA* |
| 22 | 41281613 | 41783081 | *EP300* |
| 22 | 46339498 | 47140067 | *PPARA* |
| X | 12763286 | 13225286 | *TMSB4X* |

**Supplementary Table 3: SPrime-inferred archaic introgressed segments that overlap with sequenced regions of HAA-related genes (YRI as outgroup)**

| Chromosome | Position Range (Hg19) | Target Gene | Overlapping Genes | Location to the Target Gene | Segment Label | Match Rate with Altai Neanderthal (%) | Match Rate with Altai Denisovan (%) |
|---|---|---|---|---|---|---|---|
| 1 | 231739562-231873278 | EGLN1 | ***DISC1***, *LINC00582, TSNAX-DISC1* | Downstream | EGLN1-dA | 75.86 | 24.14 |
| 2 | 46298751-46458516 | EPAS1 | ***PRKCE*** | Upstream | EPAS1-uA | 13.51 | 83.78 |
| 2 | 46657114-46808047 | EPAS1 | ***TMEM247, ATP6V1E2, RHOQ***, *RP11-417F21.1* | Downstream | EPAS1-dA | 22.53 | 46.48 |
| 9 | 119023930-119113818 | PAPPA | -- | Core | PAPPA-cA | 82.14 | 28.57 |
| 9 | 119137972-119470221 | PAPPA | ***ASTN2, TRIM32, AL137024.1*** | Downstream | PAPPA-dA | 48.31 | 14.41 |
| 12 | 33736191-34854345 | SYT10 | ***ALG10***, *RP13-359K18.1* | Downstream | SYT10-dA | 76.18 | 4.22 |
| 12 | 52432776-52626674 | ACVRL1 | ***NR4A1, OR7E47P, ATG101***, *RP11-1100L3.7*, ***KRT80*** | Downstream | ACVRL1-dA | 75.81 | 9.68 |
| 16 | 89659406-90188467 | FANCA | ***CPNE7, CDK10, DPEP1, CHMP1A, SPATA2L, SPIRE2, MC1R, TCF25, CENPBD1, DBNDD1, SPATA33, AC092143.1***, *VPS9D1-AS1*, ***ZNF276, DEF8, TUBB3***, *RP11-566K11.5, AFG3L1P* | Upstream, Core, Downstream | FANCA-cA | 76.68 | 51.21 |
| 22 | 46408289-46480570 | PPARA | *CITF22-92A6.1, LINC00899*, ***PRR34***, *RP6-109B7.5, PRR34-AS1, RP6-109B7.2, RP6-109B7.4, MIRLET7BHG* | Upstream | PPARA-uA | 9.09 | 70.45 |

*The above table shows the Denisovan introgressed segments in Tibetans that overlap with high altitude adaptation (HAA) candidate gene regions (segments inferred by SPrime). Each row represents a unique segment. From left to the right, each column denotes the chromosome of the segment, the genomic coordinate ranges (hg19), the nearby core HAA gene, other overlapped genes (protein-coding genes in bold), the relative location to the nearby HAA gene, the label of the segment (with "-u/c/d" corresponding to upstream, core, and downstream, and "A" corresponding to SPrime outgroup being Yorubans), match rate to Altai Neanderthal, and match rate to Altai Denisovan.*

**Supplementary Table 4: Significant high scoring biological pathways ranked by subnetwork score (*p*-value < 0.05)**

| Pathway | Network size | Subnetwork size | Subnetwork score | *p*-value | Subnetwork genes |
|---|---|---|---|---|---|
| p73 transcription factor network | 76 | 5 | 10.29656462 | 0.018240343 | *BUB3 CLCA2 SIRT1 TP73 WWOX* |
| TNF receptor signaling pathway | 34 | 3 | 10.08066121 | 0.019313305 | *MAP2K3 TRADD TXN* |
| p38 MAPK signaling pathway | 22 | 3 | 10.08066121 | 0.019313305 | *MAP2K3 MAP3K5 TXN* |
| Insulin Pathway | 42 | 2 | 8.891866245 | 0.0472103 | *F2RL2 RHOQ* |
| Insulin-mediated glucose transport | 17 | 2 | 8.891866245 | 0.0472103 | *RHOQ VAMP2* |

*This table shows the biological pathways (NCI database) that are significant (p-value < 0.05) for being enriched with archaic introgressed alleles and are under positive selection in Tibetan population. The archaic alleles used in this analysis include all diagnostic SNPs identified from SPrime. Each row in this table represents a pathway, and each column from left to right represents the pathway's name, the total number of genes included in the pathway (network size), the number of genes involved in positive selection for archaic alleles (subnetwork size), the score assigned from the high scoring subnetwork (HSS) analysis, the p-value of the pathway, and the names of the genes that are under positive selection (subnetwork genes, total number corresponding to the subnetwork size).*

**Supplementary Table 5: Time Estimates on Denisovan admixture, Tibetan-Han Chinese Divergence, and *EPAS1* selection from this and other studies.**

| **Date (ka)** | | **Summary Statistics/Methods** | **Type of Data** | **Reference** |
|---|---|---|---|---|
| **Denisovan Admixture Time in Asia** | ***EPAS1* Selection Time** | | | |
| 43.53 [60.00-15.70] **estimate for introgression in East Asia* | 12.30 [50.00-1.93] | Distribution of introgressed tract length /ABC | *EPAS1* gene sequence (Tibetans) | This study |
| 45.7 (31.9-60.7) **Shared with Papuans* | - | Distribution of tract length/Maximum likelihood | Whole genome sequence (Papuans) | Jacobs *et al*. 2019[6] |
| 32-12 | 12.30 (28-7) | Allele frequency, haplotype homozygosity/ Maximum likelihood | Whole genome sequence (Tibetans) | Hu *et al*. 2017[7] |
| 62-38 | - | Pairwise nucleotide difference/TMRCA | Whole genome sequence (Tibetans) | Lu *et al*. 2016[8] |
| 44-54 | - | Decay of linkage disequilibrium/ Maximum likelihood | Whole genome sequence (Papuans) | Sankararaman *et al*. 2016[9] |
| - | 12.80 (12.07–14.73) **selection not on Denisovan allele* | Extended haplotype homozygosity | Microarray data of *EPAS1* (Tibetans) | Lou at al. 2015[10] |
| - | 18.25 (17.57-18.93) | Extended haplotype homozygosity | Re-sequencing of *EPAS1* (Tibetans) | Peng *et al*. 2010[11] |

*In this table, 95% confidence intervals (CI) estimated from previous studies are shown in the parentheses. 95% credible intervals estimated from ABC in this study are shown in brackets. Lu et al. 2016 estimated the Time to the Most Recent Common Ancestor (TMRCA) as a proxy of admixture time, assuming the Denisovan and Tibetan lineages coalesce right before the introgression time. Jacobs et al. 2019 inferred 3 pulses of Denisovan admixture in Asia, with one introgressed into the Papuan lineage at 29.8 ka (95% CI 14.4–50.4), and one into the shared lineage of Papuans and other Asians at 45.7 ka (95% CI 31.9–60.7) respectively. That work did not provide an estimate on the East Asian-specific introgression time.*
